# Overcoming subtype-specific immune resistance in small cell lung cancer through RNA-based immunotherapy

**DOI:** 10.64898/2026.11.17.767319

**Authors:** Maik Schork, Tommaso Torcellan, Ioanna Tsoukala, Aniello Federico, Jennifer V. Gerbracht, Elif Diken, Özlem Akilli, Julia Zimmermann, Kimberly Wohde, Ines Beulshausen, Martin Suchan, Kevin Berg, Krutika Khinvasara, Filippo Beleggia, Franziska Lang, Barbara Schrörs, H. Christian Reinhardt, Laura Kolb, Ugur Sahin, Mustafa Diken, Ayline Kübler

## Abstract

Small cell lung cancer (SCLC) comprises neuroendocrine (NE) and non-neuroendocrine (non-NE) subtypes with distinct immune landscapes and responses to therapy. Here, we established murine experimental metastases models to investigate RNA vaccination across SCLC subtypes. These exhibited distinct organotrophic properties and differ in both the magnitude and kinetics of interferon-induced MHC-I upregulation. Vaccination controlled non-NE tumors but provided limited survival benefit against NE tumors established in the lung or liver. Single-nucleus RNA sequencing and chromatin accessibility profiling revealed inflammation-associated programs following vaccination shared between organs. To overcome the limited survival benefit of vaccination in NE SCLC, we combined RNA vaccination with liver-targeted RNA-encoded cytokines. This combination markedly prolonged survival of NE tumor-bearing mice and enhanced inflammatory immune activity within the tumor microenvironment. Together, these findings identify vaccination-associated inflammatory programs of potential clinical relevance and warrant further investigation of cytokine combination therapy as strategy to improve immunotherapy in NE SCLC.

## Introduction

SCLC is an aggressive pulmonary neuroendocrine malignancy characterized by rapid progression, early metastasis, and a 5-year survival rate below 15%^1^. Standard treatment combines platinum-based chemotherapy with immune-checkpoint blockade or radiotherapy^2^. The more recent approval of tarlatamab, a DLL3-directed CD3 T-cell engager, highlights the potential of T-cell redirection while underscoring the need for more effective and durable immunotherapeutic strategies^3^. SCLC is highly heterogeneous and comprises transcriptional states defined by lineage-associated regulators broadly classified into NE and non-NE subtypes^4–6^. ASCL1- and NEUROD1-driven tumors constitute the classical NE SCLC-A and SCLC-N subsets, respectively, whereas POU2F3-high SCLC-P tumors are rare non-NE tuft cell-like neoplasms^7,8^. A fourth, YAP1-associated program is linked to loss of canonical NE features and increased cellular plasticity; although its precise classification remains debated, it is generally considered part of the non-NE spectrum^6,9,10^.

These transcriptional states are functionally linked to distinct immune phenotypes^6,11,12^: while SCLC is generally immune-cold despite its high mutational burden, with limited immune infiltration and impaired antigen presentation, non-NE tumors typically show greater immune-cell presence than NE tumors^6,11–14^. Consistent with this, retrospective clinical analyses suggest that responses to immune checkpoint blockade are enriched in patients with non-NE SCLC^6^. However, recent work has refined this view by identifying an inflamed SCLC state characterized by immune-active features across both NE and non-NE tumors^10^. These programs are dynamic and can coexist within individual tumors, but reductionist models capturing dominant states provide a tractable framework to study state-specific tumor–immune interactions^6,10,15^.

RNA-based immunotherapies offer a flexible approach to enhance anti-tumor immunity^16^. In mutationally rich tumors, neoantigen-targeted RNA vaccination increases T-cell based antitumoral effects by inducing neoantigen-specific T cells in both preclinical and clinical settings^17^. Locally delivered RNA-encoded cytokines can further reshape the TME by promoting innate and adaptive immune activation^18,19^.

Here, we established syngeneic allograft models capturing key immune features of NE and non-NE SCLC to define subtype-specific tumor-immune interactions and responses to RNA-based immunotherapy. Notably, model-antigen expression was sufficient to trigger rejection in non-NE SCLC, whereas NE SCLC showed limited T-cell recognition at steady state. Interferon (IFN)-mediated MHC-I upregulation on NE tumor cells increased vaccination efficacy in a prophylactic setting. Therapeutic neoantigen-targeted RNA-LPX vaccination prolonged survival in non-NE tumor-bearing mice but was less effective in NE SCLC, irrespective of metastatic site. Single-nucleus RNA and ATAC sequencing after vaccination identified IFN-response, inflammatory, and MHC-I presentation programs shared across lung and liver metastasis models of NE SCLC.

To improve survival in NE SCLC, we combined neoantigen-targeted RNA vaccination with mRNA-encoded IFN-α and IL-15sushi, designed to enhance innate and adaptive antitumor were immunity. This combination significantly prolonged survival in NE SCLC-bearing mice.

Together, our findings show that tumor-cell state, rather than metastatic site, is a major determinant of immunotherapy responsiveness in SCLC. Rational combination of RNA-based vaccination and cytokine delivery can reprogram the TME and overcome immune resistance.

## Results

### Establishment of SCLC cell lines recapitulating features of immune infiltrated non-NE and immune-excluded NE states

To uncover therapeutic vulnerabilities in distinct SCLC states, we established murine tumor models using SCLC cell lines that recapitulated immune-hot and immune-cold features associated with NE and non-NE SCLC subtypes, respectively. Briefly, the tumor suppressor genes *Rb1* and *Tp53* were conditionally inactivated in the lungs of *Rb1^fl/fl^;Tp53^fl/fl^* (RP)-C57BL/6 mice through intratracheal instillation of Cre-expressing viral vectors, leading to the development of lung tumors used to generate cell lines *ex vivo*, as previously described^20^. Cell lines were derived from tumors and cultured as spheroids (3D) or as adherent cell (2D) (Extended Data Fig. 1A), resulting in distinct gene expression profiles (Extended Data Fig. 1B). 3D culture favored NE-associated markers (*Ascl1*, *Chga*, *Chgb*), whereas 2D culture enriched non-NE markers (*Yap1*, *Rest*, *Vim*) (Fig. 1A, Extended Data Fig. 1C), consistent with SCLC-A and SCLC-Y subtype identities, respectively (Fig. 1A). SCLC-Y1 and SCLC-A1/A3 were selected for subsequent experiments.

**Figure 1:**
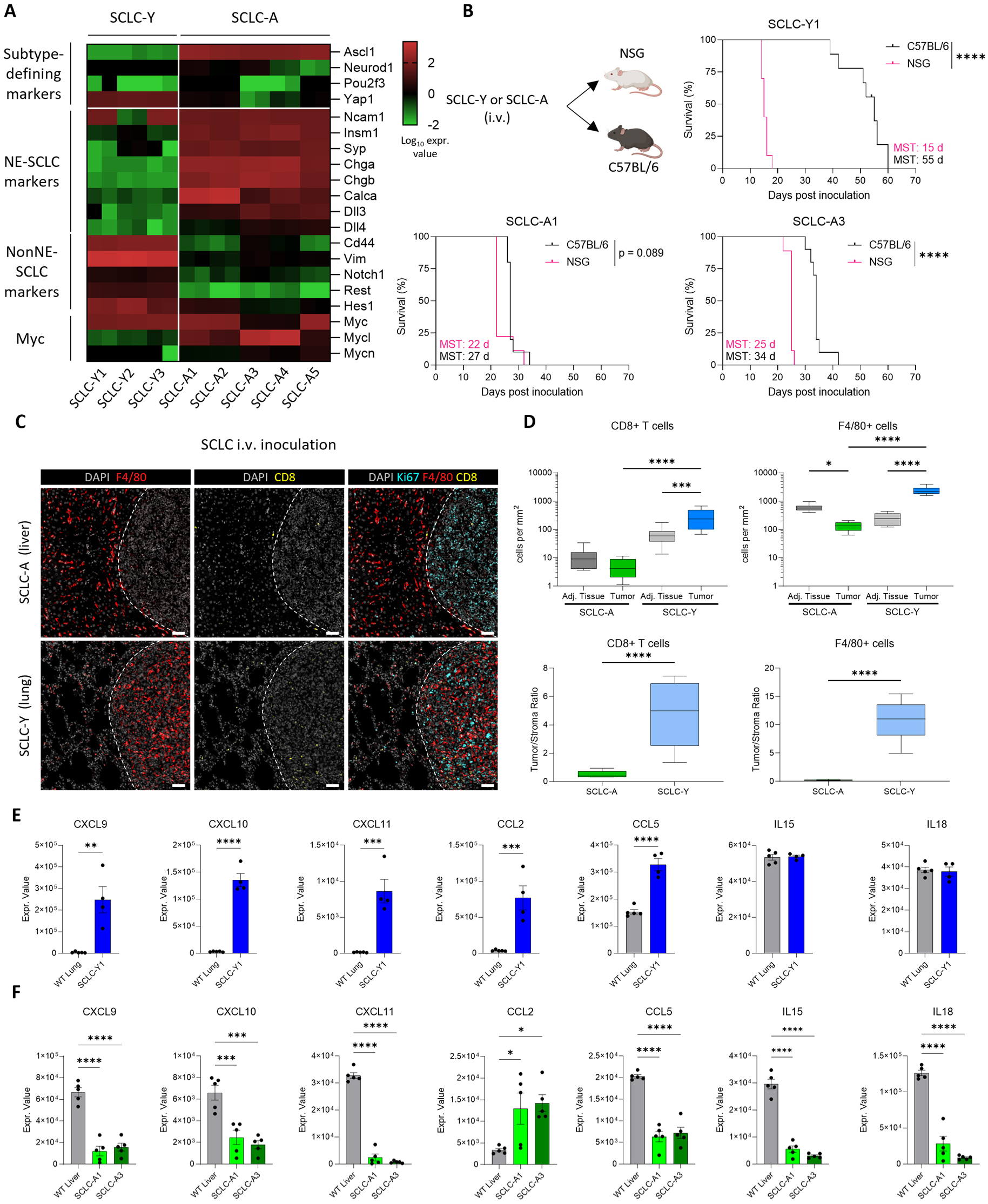
Ascl1-high and Yap1-high SCLC tumors display distinct engraftment, immune infiltration and chemokine profiles. (A) Gene expression analysis of subtype-defining, neuroendocrine and non-neuroendocrine marker genes in murine SCLC cell lines via bulk RNA sequencing. Shown are technical duplicates as log10 fold expression values. (B) C57BL/6 or NSG mice (n = 10) were intravenously injected with 5 x 10^6^ tumor cells. Mice were monitored until termination criteria were reached. Significance for survival was determined via Mantel-Cox logrank test. (**C, D**) Tumor-bearing mice (n = 5) were histologically analyzed at the endpoint for CD8 and F4/80 by immunohistochemistry. (C) Representative staining of tumor nodules (DAPI white, CD8 green, F4/80 red), tumor tissue is indicated with dashed line, scale bar 50 µm. (D) Quantification of CD8^+^ or F4/80^+^ cells in tumor or adjacent liver (SCLC-A) or lung (SCLC-Y) tissue. Significance was determined by one-way ANOVA with posthoc Tukey’s multiple comparison test (upper graphs) or t-test (lower graphs). (**E, F**) Cytokine and chemokine expression was quantified by fluidigm q-RT-PCR in livers (SCLC-A) and lungs (SCLC-Y) of tumor-bearing mice (n = 5 for SCLC-A; n = 4 for SCLC-Y) at study endpoint and organs of naïve mice as control (n = 5). Shown expression values were calculated by ΔΔCt normalized to housekeeping genes. Significance was determined by one-way ANOVA with posthoc Tukey’s multiple comparison test. Significance levels for full figure are displayed as following: p < 0.05 = *; p < 0.01 = **; p < 0.001 = ***; p < 0.0001 = ****

Engraftment of both NE SCLC-A and non-NE SCLC-Y tumors upon intravenous (i.v.) cell line injection was delayed in immunocompetent mice compared to immunodeficient mice, suggesting an important role for immune recognition, particularly in restraining SCLC-Y tumor development (Fig. 1B). Histological analysis confirmed that subtype-defining markers continued to be expressed in the engrafted tumors (Extended Data Fig. 1D).

Interestingly, SCLC-Y and SCLC-A tumors displayed organotropic properties, as SCLC-Y tumors preferentially engrafted in the lungs, whereas SCLC-A tumors predominantly developed in the liver as indicated by increased organ weight (Extended Data Fig. 1E) and release of liver damage-associated enzymes (Extended Data Fig. 1F) as observed in patients^21^. Immunohistochemistry revealed increased F4/80⁺ macrophage and CD8⁺ T-cell infiltration in SCLC-Y tumors relative to adjacent healthy tissue, whereas SCLC-A tumors exhibited an immune-excluded phenotype (Fig. 1C, D), consistent with clinical observations^11^. To assess whether immune infiltration was subtype-rather than site-dependent, SCLC-A cells were additionally inoculated intrapulmonarily, where tumors retained the immune-deserted phenotype observed in liver-engrafted tumors (Extended Data Fig. 1G, H).

Consistently, SCLC-Y lung tumors showed increased cytokine and chemokine expression relative to healthy tissue as determined by q-RT-PCR, whereas for SCLC-A this inflammatory signature was largely absent (Fig. 1E, F). Together, these data indicate that SCLC-Y1 and SCLC-A1/A3 recapitulate distinct immune states characteristic of non-NE and NE SCLC.

### SCLC-A and SCLC-Y tumors differ in T-cell recognition

To determine how expression of a model neoantigen influences immune-mediated tumor rejection, we engineered SCLC-A3 and SCLC-Y1 tumor cell lines for stable and comparable expression of the previously described mutant Rpl18 and Kif18b as model neoantigens^22,23^ together with GFP separated by a P2A sequence. Naïve C57BL/6 mice were injected i.v. with transduced or wildtype cell lines (Fig 2A). In contrast to their wildtype counterparts, model antigen-expressing SCLC-Y tumors were efficiently rejected in the absence of therapeutic intervention, whereas SCLC-A injection resulted in rapid and lethal tumor progression that was only minimally affected by model antigen expression (Fig. 2B).

**Figure 2:**
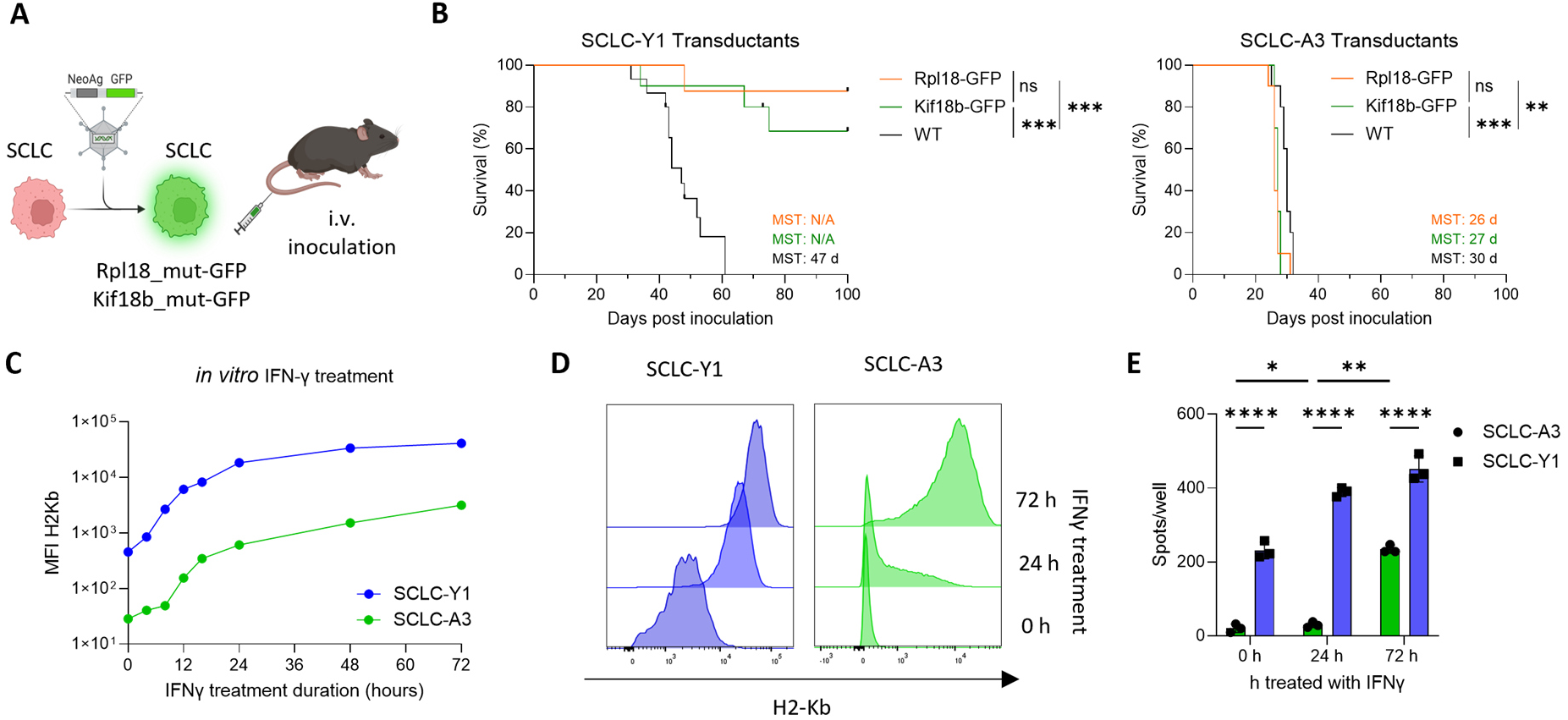
Distinct SCLC subtype responses to neoantigen expression and subtype-specific IFN-γ–induced MHC-I upregulation. (**A**) SCLC cell lines were retrovirally transduced with model antigens Rpl18-GFP or Kif18b-GFP (Created in BioRender. https://BioRender.com/bm44j6p) and (**B**) intravenously injected into C57BL/6 mice (n = 10). Mice injected with non-transduced cell lines served as control. Mice were monitored until termination criteria were reached. Significance for survival was determined via Mantel-Cox logrank test. (**C**) Tumor cell lines were treated with 50 ng/mL IFN-γ for the indicated time periods, and the mean fluorescent intensity (MFI) of H-2Kb was measured **(D)**. Rpl18-GFP transduced cell lines were pre-treated with 50 ng/ml IFN-γ for 0, 24 and 72 h and analyzed for H-2Kb expression before co-culture. (**E**) Splenocytes isolated from mice vaccinated against the Rpl18 neoantigen were co-cultured with Rpl18 neoantigen-transduced SCLC cell lines pre-treated with IFN-γ for the indicated periods of time. Shown are IFN-γ spot counts per well in technical replicates (n = 3). Significance was determined by two-way ANOVA with Tukey’s post hoc test for multiple comparisons. Shown are selected significance comparisons. p < 0.05 = *; p < 0.01 = **; p < 0.0001 = ****

To assess whether tumor rejection was driven by high neoantigen expression, we sorted GFP-low SCLC-Y1 cells, which likewise failed to fully engraft, except for one lung containing small GFP-negative tumor islands, potentially reflecting transgene loss (Extended Data Fig. 2A-C). Thus, tumor rejection was not simply attributable to artificially high model antigen expression. As reduced antigen presentation can impair T-cell recognition and promote immune escape in SCLC^14^, we examined expression of key components of the antigen presentation machinery. SCLC-A cell lines generally exhibited lower expression of *Tap1*, *Tap2* and *Nlrc5* transcripts than SCLC-Y (Extended Data Fig. 2D). In the absence of stimulation, SCLC-Y1 cell lines expressed moderate and SCLC-A cell lines low levels of MHC-I protein (Fig. 2C, D). However, upon 24 h IFN-γ treatment, MHC-I expression was readily induced in SCLC-Y, whereas SCLC-A displayed delayed and attenuated MHC-I induction (Fig. 2C, D), depending rather on duration of the IFN-γ exposure than dose (Extended Data Fig. 2E). To investigate whether these differences in MHC-I induction translate into differential T-cell recognition, we performed an *in vitro* IFN-γ ELISpot assay using splenocytes from Rpl18-neoantigen vaccinated mice. While SCLC-Y1 Rpl18-GFP tumor cells elicited T-cell responses even without prior IFN-γ stimulation, SCLC-A3 Rpl18-GFP tumor cells required 72 h of IFN-γ pretreatment to elicit substantial responses.

Together, these findings suggest that the delayed and reduced MHC-I upregulation in NE SCLC may limit its recognition by antigen-specific T cells.

### SCLC subtypes differ in their susceptibility to neoantigen vaccination

Although expression of strong model neoantigen revealed marked differences in immune recognition between SCLC-A and SCLC-Y tumors, it remained unclear whether naturally occurring tumor neoantigens could be effectively targeted by therapeutic RNA-LPX vaccination and would elicit similar subtype-specific responses. To identify tumor-specific mutations for immunogenicity testing, we performed whole-exome and RNA sequencing of SCLC cell lines. Shared, as well as cell-lines-specific mutations were identified (Extended Data Fig. 3A), selected based on MHC binding ability and variant allele expression^24^ and incorporated into neoantigen-candidate encoding RNA-vaccine constructs (Fig 3A, Extended Data Fig. 3B). Immunogenicity testing using splenocytes from mice immunized with the vaccine constructs revealed several immunogenic SNV- and insertion/deletion (InDels)-derived neoantigens (Extended Data Fig. 3C, D). A further round of immunogenicity testing revealed that the identified neoantigens were predominantly CD4-restricted (Fig 3B, Extended Data Fig. 3E). Notably, NE and non-NE cell lines displayed comparable mutational burden (Extended Data Fig. 3A), suggesting that SCLC-A and SCLC-Y tumors did not exhibit major differences in predicted SNV-derived neoantigen quantity between SCLC-A and SCLC-Y (Extended Data Fig. 3F).

**Figure 3:**
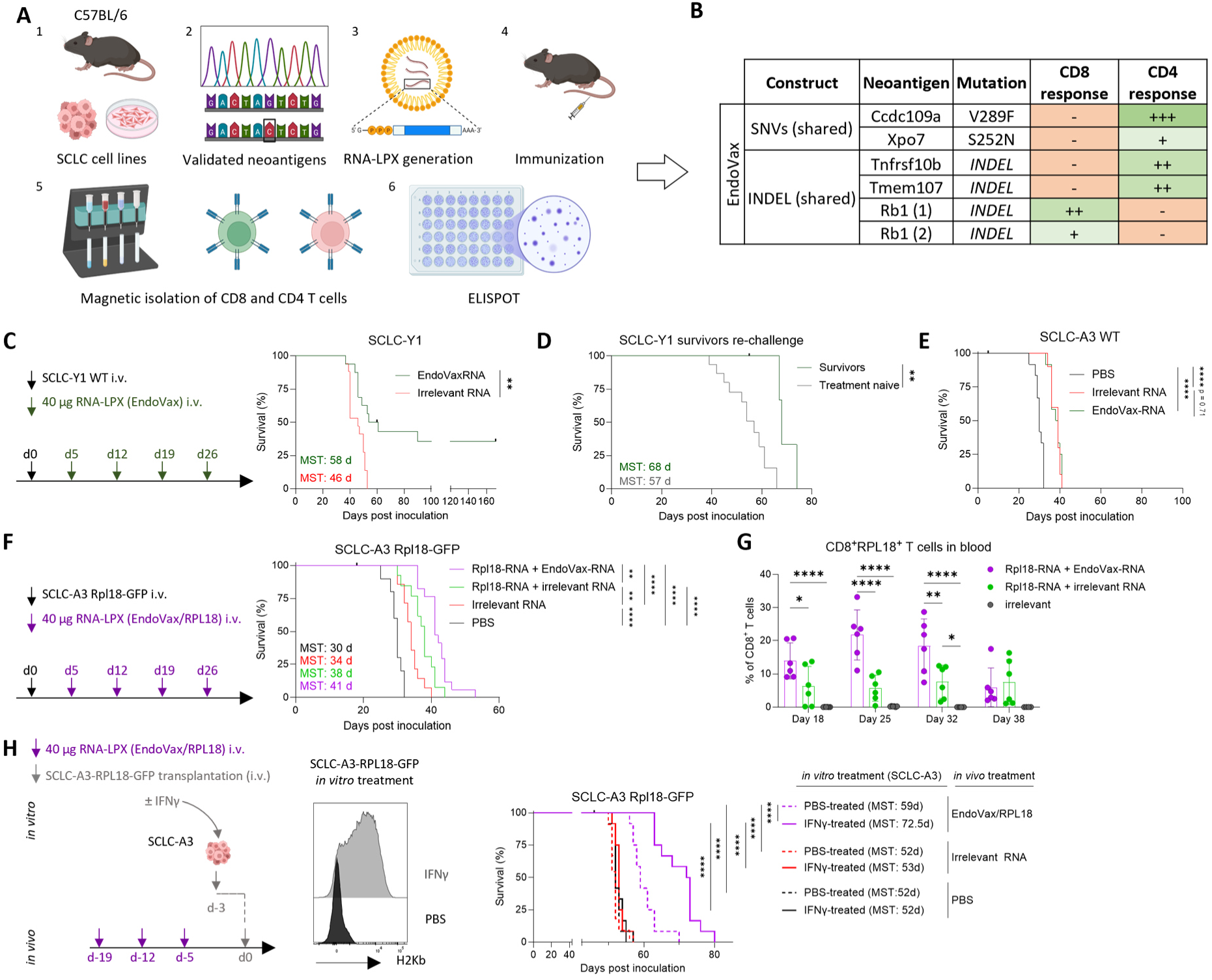
SCLC can be targeted by tumor-specific antigen RNA-LPX vaccination. (**A**) Schematic overview of the tumor-specific validation pipeline and the resulting validated shared neoantigens, comprising single nucleotide variants (SNVs) and insertions/deletions (InDels), used for subsequent vaccination studies (“EndoVax”). Created in BioRender. https://BioRender.com/ritxlvo (**B**) Validated immunogenic tumor-specific antigens. (**C**) C57BL/6 (n = 15 - 16 mice were intravenously inoculated with SCLC-Y1 cells and treated with RNA-LPX vaccination at the indicated time points; (**D**) Survivor mice from the EndoVax group in (**C**) (n = 3) and naïve C57BL/6 mice (n = 15) were re-challenged with SCLC-Y1 cells i.v.. (**E**) C57BL/6 (n = 10 - 12) were analyzed after injection i.v. with SCLC-A3 cells; and treated as indicated in C. (**F**) C57BL/6 (n = 14 - 17; PBS group was added for visualization and not part of the experiment.) were i.v. injected with SCLC-A3-GFP-Rpl18 cells and vaccinated as indicated. Shown is survival according to termination criteria. (**G**) Flow cytometric analysis of peripheral blood Rpl18-antigen specific CD8^+^ T cells from mice (n = 6) shown in (**F**) at the indicated time points. Significance was determined by two-way ANOVA with Tukey’s post-hoc test for multiple comparisons. (**H**) C57BL/6 were prophylactically vaccinated with Rpl18 and Endovax encoding RNA-LPX, irrelevant RNA-LPX or PBS as indicated. Mice were either injected with untreated SCLC-A3-Rpl18 cells (n = 11 - 12), SCLC-A3-Rpl18 cells *in vitro* pre-treated for 72 hours with 50 ng/mL IFN-γ. Shown is survival according to termination criteria, significance was determined via Mantel-Cox logrank test with comparison between each EndoVax/Rpl18 subgroup and among the corresponding *in vitro* pretreatment conditions across EndoVax/Rpl18 (*in vivo*) and PBS (*in vivo*) treated mice. Significance for all other survival curves was determined via Mantel-Cox logrank test. For all indicated significancy levels p < 0.05 = *; p < 0.01 = **; p < 0.0001 = ****

Validated neoantigens (Fig. 3B), derived from shared single nucleotide variants (SNVs) and InDels were then incorporated into a fused construct (Endovax) used for therapeutic RNA-LPX vaccination in SCLC-Y1, SCLC-A1 and SCLC-A3 tumors. Vaccination with Endovax significantly prolonged survival in SCLC-Y1-bearing mice, with a subset achieving durable long-term survival (Fig. 3C). Upon tumor re-challenge, surviving mice exhibited prolonged survival compared to naïve controls (Fig. 3D). In contrast, neither SCLC-A3 (Fig. 3E) nor SCLC-A1 (Extended Data Fig. 3G) responded to shared or unique neoantigen-encoding RNA-LPX vaccination. In Rpl18-neoantigen overexpressing SCLC-A3, therapeutic vaccination targeting the antigen induced limited anti-tumor activity, which was enhanced by combination with the endogenous neoantigen-coding construct EndoVax (Fig. 3F). Frequencies of Rpl18-tetramer positive CD8^+^ T cells were increased upon Rpl18 and EndoVax RNA combination, compared to Rpl18-neoantigen RNA-LPX vaccination alone (Fig. 3G), which may reflect CD4 T-cell help boosting Rpl18-specific CD8 T-cell responses. The combination of Rpl18 and EndoVax-based vaccination was also efficacious in a prophylactic vaccination setting, and vaccination efficacy was markedly improved when tumor cells were pre-stimulated with IFN-γ for 72 hours to upregulate MHC-I expression prior to injection (Fig. 3H).

Together, these results highlight differential subtype sensitivity to tumor antigen vaccination: Endogenous neoantigens were sufficient to induce durable anti-tumor immunity against SCLC-Y1, whereas therapeutic benefit in SCLC-A3 required vaccination against a strong model antigen and was further enhanced by combinatorial neoantigen vaccination and MHC-I upregulation on tumor cells. As SCLC-A3 consistently recapitulated key features of immune-resistant NE SCLC while exhibiting the potential to be sensitized to neoantigen vaccination, it was selectedas a NE model for subsequent mechanistic and therapeutic studies. SCLC-Y1 and SCLC-A3 are hereafter referred to as SCLC-Y and SCLC-A, respectively unless specified otherwise.

### RNA-LPX vaccination mediates comparable anti-tumoral efficacy against SCLC-A tumors in the lung and liver, along with coordinated transcriptional and epigenetic TME remodeling

As described above, SCLC-A tumors showed higher resistance to immunotherapy treatment, consistent with higher responsiveness of the non-NE subtype to checkpoint inhibition in clinical analyses^6^. We therefore focused further efforts on characterizing immune resistance mechanisms of SCLC-A, to inform on strategies to enhance the efficacy of immunotherapy treatment. To determine whether the limited responsiveness of SCLC-A to RNA-LPX reflected intrinsic subtype characteristics^6^ or was imposed by the liver microenvironment, we established SCLC-A tumors in the liver or lung via i.v. or intrapulmonary (i.pulm.) injection, respectively. Interestingly, Rpl18/EndoVax RNA-LPX vaccination conferred a comparable therapeutic benefit, as measured by overall survival, irrespective of tumor location (Fig. 4A, B), suggesting that differential vaccine responsiveness is primarily associated with SCLC subtype rather than host tissue.

**Figure 4:**
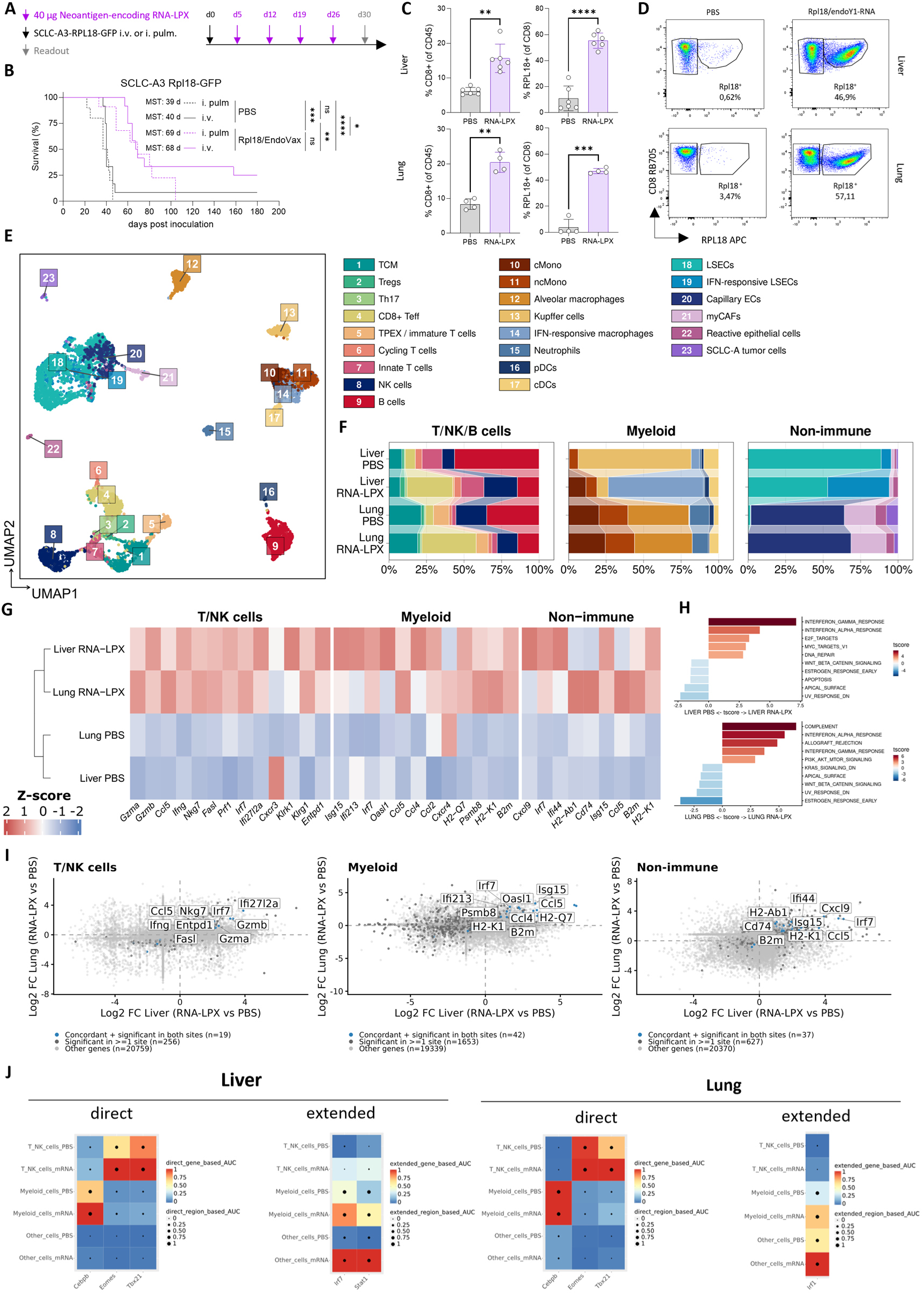
Vaccination is effective for SCLC-A tumor nodules in lung and liver and remodels the TME. **(A)** C57BL/6 mice (n = 10 - 12) i.v. inoculated with SCLC-A3-GFP-Rpl18 tumor cells to yield metastases in the liver or injected intrapulmonarily to generate tumors in the lung and treated with Rpl18/Endovax RNA-LPX vaccination as indicated. At d 20 n = 4 – 6 mice per group were sacrificed for flow cytometric and snRNAseq and ATAC analysis. (**B**) Survival analysis according to termination criteria, significance for survival was determined via Mantel-Cox logrank test. (**C-D**) Flow cytometric analysis of CD8^+^ T cells in tumor-bearing lungs or livers (n = 4 -6). (**E**) Multiome (snRNASeq and snATACSeq) dataset displayed as UMAP. Clusters were identified according to specific gene sets referenced in Extended Data Fig. 6B. (**F**) Cluster frequencies based on tumor-bearing organ (liver, lung) **and** treamtent (PBS, RNA-LPX) conditions. (**G**) Hierarchically clustered heatmap of selected differentially expressed genes across T/NK, myeloid and non-immune compartments. (**H**) Waterfall plots showing hallmark pathways (ranked by t-scores) comparing PBS and RNA-LPX treatment in liver and lung CD8 effector T cells. (**I**) Scatter plots showing concordance of treatment-associated gene expression changes between liver and lung across cell compartments. Highlighted genes have been found significantly and concordantly regulated at both sites, as also showing a concomitant enrichment in the ATAC analyses. (**J**) SCENIC+ eRegulon analysis in liver and lung. Direct and extended eRegulons are shown separately. Color indicates gene-based eRegulon activity and dot size indicates region-based eRegulon activity. Only positively regulated eRegulons are shown.

RNA-LPX vaccination strongly increased Rpl18-specific CD8⁺ T cells in both organs, with enhanced proliferation and effector differentiation (Ki67, KLRG1) as measured by flow cytometry. CD8^+^ T cells in liver metastases showed higher CD69 and PD-1 expression than lung tumors, indicating organ-specific differences in T-cell activation (Fig. 4C, D; Extended Data Figs. 4A, B and 5A, B). Vaccination also increased macrophage activation markers CD69, CD86 and CD40 across both sites albeit to varying extent (Extended Data Fig. 5C). To further characterize how RNA-LPX vaccination shapes the TME in SCLC-A tumor-bearing lungs or metastasis-bearing livers, we performed single-nucleus multiome RNA (snRNAseq) and ATAC (snATACseq) sequencing.

The integrated multiome analyses identified 23 distinct immune and non-immune cell clusters, (Fig. 4E, Extended Data Fig. 6A, B). Consistent with flow cytometry, RNA-LPX vaccination increased the relative abundance of CD8^+^ effector T cells in both organs (Fig. 4F). In metastasis-bearing livers, RNA-LPX vaccination was associated with lower relative abundance of Kupffer cells and higher relative abundance of IFN-responsive macrophages (Fig. 4F) expressing strong IFN-response signatures and co-stimulatory molecules (*Cd80, Cd86*; Extended Data Fig. 6B). Among the non-immune cells, RNA-LPX-vaccinated liver tumors displayed a markedly higher proportion of IFN-responsive liver sinusoidal endothelial cells (LSECs) (Fig. 4F), characterized by high expression of genes such as *Cxcl9*, *Cxcl10*, *Vcam1*, and *Isg15* (Extended Data Fig. 6B). Differential gene expression analysis further revealed broad transcriptional changes following RNA-LPX vaccination. T and NK-cell populations upregulated cytotoxicity-related genes, including *Gzmb, Fasl, Prf1, Cx3cr1* and *Nkg7* in (Fig. 4G, Extended Data Fig. 6C, D), alongside reduced expression of naïve/memory-associated markers (*Sell*, *Ccr7, Il7r*) and residency-associated markers (*Cxcr6, Itga1*) consistent with a shift towards an effector phenotype (Extended Data Fig. 6C). Myeloid cells upregulated chemokines (*Ccl2,Ccl4)*, components of the antigen presentation machinery (*Psmb8, H2-K1, B2m*) and proinflammatory genes (Fig. 4G, Extended Data Fig. 6E). Some of the transcriptional changes were also observed in non-immune populations (Fig. 4G). Together, these findings revealed in both tumor sites, broad immune and stromal transcriptional remodeling following RNA-LPX vaccination, characterized by enhanced cytotoxic programs and interferon-associated responses across multiple cellular compartments (Fig. 4H, Extended Data Fig. 6F).

We next investigated the contribution of epigenetic remodeling by analyzing differently accessible regions (DARs) between treated and untreated conditions across the different cell types and anatomical sites. Significant DARs were identified predominantly among non-immune and myeloid cells in livers and, whereas in the lungs, DARs were primarily identified in T cells (Extended Data Fig. 7A). Notable gene ontology (GO) pathways linked to DARs included immune cell activation, tissue migration and immune response signaling (Extended Data Fig. 7B). To assess concordance between transcriptional and epigenetic changes, we analyzed the overlap between differentially expressed genes (DEGs) and DARs (Extended Data Fig. 7C), identifying gene expression changes driven by epigenetic reprogramming. We further compared the log-fold changes of DAR-associated DEGs between lung and liver, revealing a group of genes with concordant expression changes across organs. These included genes, including genes related to cytotoxic effector functions, interferon response and antigen presentation (Figure 4I).

To integrate transcriptional and chromatin accessibility changes, we performed SCENIC+ analysis, identifying enhancer-driven regulons (eRegulons) linking transcription factor (TF) activity to enhancer accessibility and gene expression. RNA-LPX treatment increased TF activity across T/NK, myeloid and non-immune cells in both organs (Extended Data Fig. 7D). Prominent regulons were associated with cytotoxicity and interferon responses in T/NK cells (e.g., *Tbx21, Eomes, Stat1, Irf7*), inflammatory and interferon programs in myeloid cells (e.g., *Cebpb, Fos, Nfkb1, Stat1*), and interferon signaling in non-immune cells (e.g., *Irf7, Stat1, Irf8*) (Fig. 4J). Overall, these findings indicate coordinated transcriptional and epigenetic activation of inflammatory, antigen-presentation and cytotoxicity programs following RNA-LPX treatment, consistent with a shift towards an antitumoral microenvironment.

### RNA-encoded cytokines synergize with RNA-LPX vaccination for strong antitumoral efficacy in liver metastases of SCLC-A

Collectively, our data indicated that efficient antigen presentation and interferon-responsive transcriptional programs are potential drivers of effective anti-tumor immunity in SCLC. We hypothesized that therapeutic enhancement of IFN-driven immune responses induced by liver-targeted RNA-encoded IFN-α and IL15sushi could improve RNA-LPX vaccination in SCLC-A liver metastases and further overcome immune resistance. IFN-α promotes antigen presentation, dendritic-cell activation, and T-cell priming, whereas IL-15sushi supports NK-cell and memory CD8^+^ T-cell responses^26,27^. To limit nonspecific innate activation by RNA, we used double-strand purified N1-methylpseudouridine-modified RNA.

Either RNA-encoded cytokines or RNA-LPX vaccination alone produced resulted in comparable anti-tumor effects relative to controls. In contrast, the combination markedly extended median survival to 101 days versus 34 days in PBS-treated mice (Fig. 5A, B). Spectral flow cytometry of immune cells in tumor-bearing livers at day 30 after tumor inoculation and unsupervised clustering enabled the unbiased identification of distinct cell subsets based on their phenotypic signatures, involving several CD8^+^, CD4^+^ T cell as well as ILC/NK-cell clusters (Fig. 5C, Extended data Fig. 8A, B). Comparative analysis of cluster frequencies revealed treatment-specific effects (Fig. 5D) RNA-LPX vaccination strongly expanded Rpl18-specific CD8^+^ T cells (Fig. 5D, E), including terminal effector (CD44^+^ CX3CR1^+^ KLRG1^+^, cluster 4) and memory precursor/early effector (CD44^+^ KLRG1^-^CD127^+/-^, cluster 8) populations (Fig. 5F; Extended Data Fig. 8C). Although combination therapy improved survival, reduced frequencies of Rpl18-specific clusters were detected (Fig. 5D-F). Instead, it simultaneously enriched Rpl18-negative CD8^+^ T-cell subpopulations (Fig. 5D): cluster 6, a KLRG1^+^ CX3CR1^+^ terminal effector subset induced by vaccination (Fig. 5G), and cluster 11, a CD69^+^ Eomes^+^ population lacking PD-1 and TOX consistent with a highly activated or a tissue resident memory phenotype, preferentially expanded by cytokine RNA (Fig. 5H-I; Extended Data Fig. 8). Thus, vaccination and cytokine treatment reshape distinct cytotoxic cell compartments, suggesting that the therapeutic benefit reflects broader immune remodeling rather than simply greater expansion of vaccine-specific T cells.

**Figure 5:**
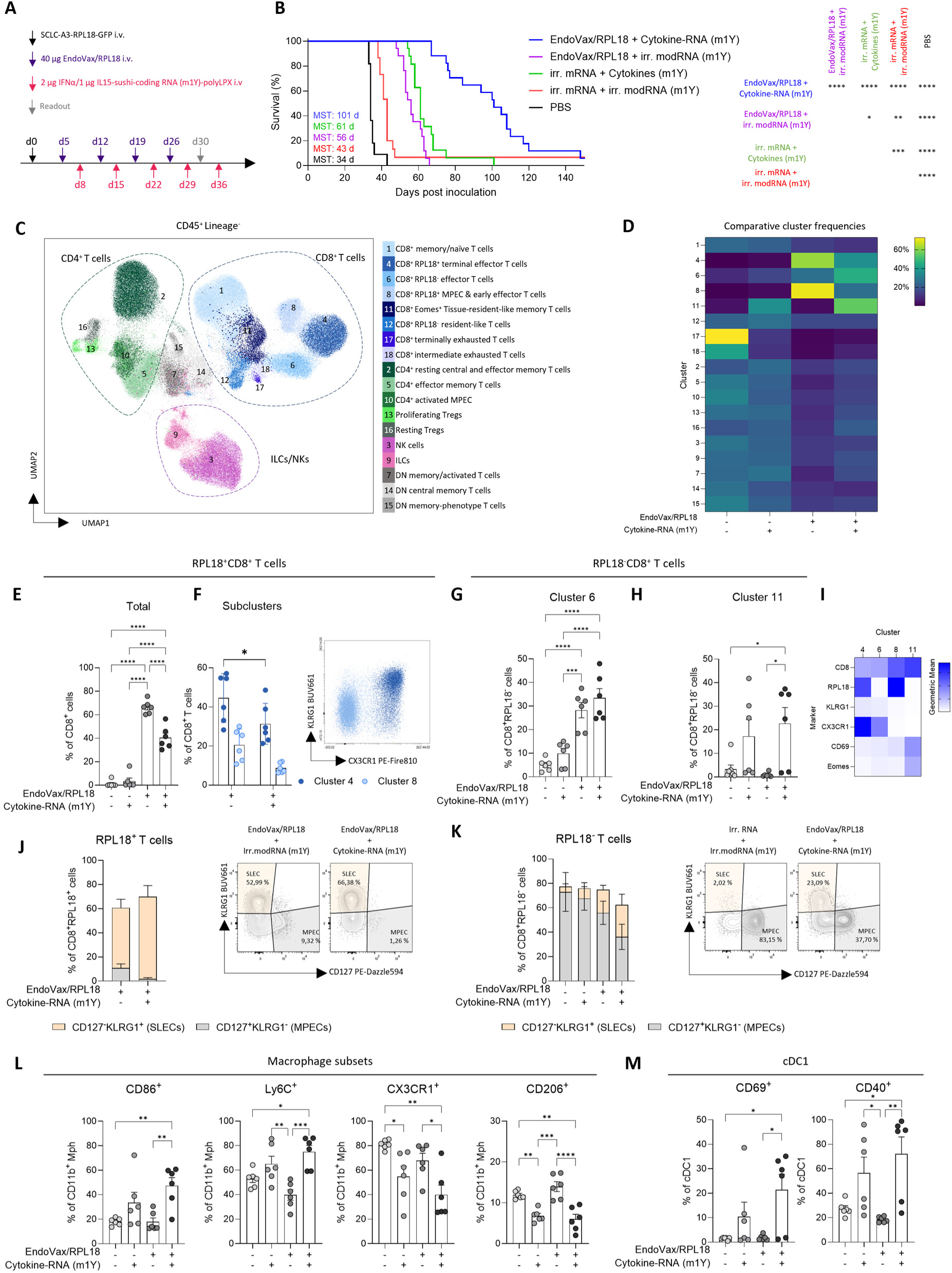
Liver-targeted RNA-encoded IFN-α and IL-15 sushi synergize with RNA-LPX vaccination for treatment of SCLC-A liver metastasis. **(A)** C57BL/6 mice (n = 15 -17 and n = 11 for PBS-treated mice) were intravenously inoculated with SCLC-A3-GFP-Rpl18 tumor cells to yield metastases in the liver and intravenously treated with Rpl18/Endovax RNA-LPX vaccination and/or RNA-encoded IFN-a and IL-15 sushi in liver-targeted formulation as indicated. At d 30 n = 6 mice per group were sacrificed for flow cytometric analysis. (**B**) Survival analysis according to termination criteria, significance was calculated by Mantel-Cox logrank test. (**C**) UMAP showing unsupervised clustering of concatenated flow cytometry data by X-Shift. (**D**) Comparative cluster frequencies (%) as identified by X-shift grouped by treatment. (**E**) Frequency of Rpl18-antigen-specific CD8+ T cells as percentage of total CD8+ T cells as identified by hierarchical gating. (**F**) RPL18-specific subclusters 4 and 8 identified by X-shift as percentage of CD8^+^ T cells and a representative flow cytometry dot plot to visualize their KLRG1 and CX3CR1 expression. (**G**, **H**) Shown are subclusters 6 and 11 as identified by X-shift as percentage of CD8^+^ T cells. (**I**) Heatmap summarizing expression (gMFI) of selected markers of clusters 4, 6, 8 and 11 as identified by X-shift. (**J**) Hierarchically gated CD127^-^ KLRG1^+^ short-lived effectors (SLEC) and CD127^+^ KLRG1^-^memory precursors (MPEC) and their percentage of CD8^+^ Rpl18-antigen-specific T cells and representative flow cytometry contour plots. (**K**) Percentage of SLECs and MPECs on non-Rpl18 specific CD8^+^ T cells and representative flow cytometry contour plots. (**L, M**) selected marker expression on CD11b^+^ F4/80^+^ macrophages (L) and cDC1(M). Significance for flow cytometric analyses as indicated was performed by one-way ANOVA and post hoc Tukey’s test for multiple comparison. Significance levels are displayed as following: p < 0.05 = *; p < 0.01 = **; p < 0.001 = ***; p < 0.0001 = ****

Hierarchical gating revealed that vaccination-induced Rpl18^+^ CD8^+^ T cells were predominantly KLRG1^+^ CD127^-^short-lived effector cells (SLEC) which only marginally changed upon cytokine co-treatment (Fig. 5J), whereas Rpl18-negative CD8^+^ T cells were mainly KLRG1^-^CD127^+^ memory precursor effector cells, which were reduced upon combination treatment in favor of SLECs (Fig. 5K), RNA-LPX vaccination increased CD8^+^ T-cell and, to a lesser extent, NK-cell frequencies and improved the CD8/Treg ratio (Extended Data Fig. 9A, B). Combination therapy further enhanced NK-cell activation, measured by CD69 expression, without substantially affecting Treg proliferation or activation (Extended Data Fig. 9C, D).

Because SCLC-A cells showed low basal MHC-I expression and poor T-cell recognition (Fig. 2), we assessed whether cytokine treatment restored antigen presentation. Cytokine RNA alone significantly increased tumor-cell MHC-I MFI in vivo (Extended Data Fig. 9E), potentially increasing susceptibility to CD8^+^ T-cell-mediated killing. Finally, combination treatment produced the strongest myeloid remodeling (Fig. 5L, M; Extended Data Fig. 9F). It increased CD86^+^ and CD40^+^ macrophages and CCR2^+^ Ly6C^+^ inflammatory subsets, while reducing CX3CR1^+^, Ly6C^-^, CD206^+^, and CD115^+^ macrophages, consistent with a more inflammatory, potentially anti-tumoral phenotype (Fig. 5L; Extended Data Fig. 9G). Cytokine or combination treatment also increased CD69, CD40, and CD86 expression on cross-presenting cDC1 cells (Fig. 5M; Extended Data Fig. 9H), potentially enhancing T-cell priming.

In summary, liver-targeted RNA-encoded IFN-α and IL-15sushi synergized with RNA-LPX vaccination in SCLC-A liver metastases. Vaccination primarily expanded Rpl18-specific T-cell responses, whereas cytokine RNA promoted broader activation of antigen-specific and bystander lymphocyte populations and pronounced myeloid activation. Cytokine treatment also increased tumor-cell MHC-I expression, collectively creating a more inflammatory and immune-permissive tumor microenvironment that likely underlies the therapeutic synergy. Overall, immune-hot SCLC-Y was susceptible to RNA-LPX vaccination across a range of antigen strengths, whereas SCLC-A required high-quality, multi-antigen targeting and additional immune modulation. RNA-encoded cytokines partially overcame these barriers by increasing tumor visibility through MHC-I upregulation, broadening T-cell activation, activating NK cells, and reprogramming myeloid cells. Together, our findings support subtype- specific strategies combining rational antigen selection with cytokine RNA-based immune modulation to overcome immunotherapy resistance in SCLC-A. Our murine models recapitulate distinct immune phenotypes of human SCLC across anatomical sites, providing a translational preclinical platform to study tumor–immune interactions and evaluate novel therapies.

## Discussion

Phase III clinical trials have demonstrated only modest therapeutic benefits of immunotherapy in SCLC, despite its high mutational burden. In IMpower133, the addition of atezolizumab to chemotherapy improved median overall survival from 10.3 to 12.3 months^26^. KEYNOTE-604 reported 12.9 versus 9.7 months with pembrolizumab plus chemotherapy versus chemotherapy alone, although the difference did not meet the prespecified statistical significance threshold^27^. Similarly, CASPIAN showed improvement from 10.5 to 12.9 months with the addition of durvalumab to chemotherapy^28^. Only recently, DeLLphi-305 data demonstrated an increase from 12.7 to 21.1 months in patients treated with tarlatamab + durvalumab vs. durvalumab only without progression after initial chemoimmunotherapy.^29^ These modest gains underscore the need to address mechanisms of immunotherapy resistance in SCLC.

SCLC subtypes differ markedly in immune contexture and sensitivity to immune checkpoint inhibitors. Inflamed non-NE subtypes show greater T-cell infiltration, higher HLA expression, and may be more sensitive to immune checkpoint inhibitors than NE subtypes such as SCLC-A^6,11,30^. In contrast, NE SCLC is often characterized as immunologically “cold,” with impaired antigen presentation and limited immune infiltration. In NE-SCLC, low HLA expression, frequently caused by epigenetic silencing of antigen-presentation machinery, as well as enrichment of immunosuppressive myeloid cells are associated with poor immunotherapy response^6,10,14,15^.

To investigate subtype-specific tumor-immune interactions, we established murine SCLC models that recapitulate features of the immunologically cold and hot TMEs characteristic of NE and non-NE SCLC, respectively. We used these models to evaluate RNA-LPX vaccination and its interplay with tumor-intrinsic immune evasion. The models recapitulate the low but inducible MHC I expression typical of NE SCLC, a key barrier to effective anti-tumor immunity.

Expression of the strongly immunogenic Rpl18 neoantigen was sufficient to induce rejection of immune-hot SCLC-Y tumors, whereas immune-cold SCLC-A tumors remained resistant. Similarly, therapeutic RNA-LPX vaccination targeting newly identified endogenous tumor antigens induced durable responses in SCLC-Y but not SCLC-A. In the latter, therapeutic efficacy was observed only when vaccination against endogenous antigens, which were predominantly CD4-directed, was combined with vaccination against the strongly immunogenic CD8 neoantigen Rpl18.

These contrasting responses may partly reflect differences in antigen-presentation capacity. While SCLC-Y cells readily upregulated MHC-I following IFN-γ stimulation, SCLC-A cells required prolonged stimulation, suggesting that stronger or more sustained immune activation may be necessary to achieve effective T-cell recognition and tumor control.

The importance of tumor-cell MHC-I expression was further supported by prophylactic vaccination experiments: while complete protection was observed in other tumor models such as CT26 or B16F10^31^, prophylactic vaccination conferred only minor survival benefit in SCLC-A. Pretreatment with IFN-γ to increase tumor-cell MHC-I expression markedly improved survival after prophylactic vaccination, supporting a role of limited MHC-I expression in the resistance of SCLC-A to vaccine-elicited immunity. Beyond tumor-cell antigen presentation, our findings suggest that CD4 T-cell help may contribute to vaccination efficacy. Combining endogenous antigens with the CD8-targeted Rpl18 antigen induced higher frequencies of Rpl18-specific T cells than CD8-targeted vaccination alone, suggesting that CD4-directed priming may enhance the magnitude and quality of cytotoxic T-cell responses.

Given the preferential liver tropism in our SCLC-A cell line system, we investigated whether its limited responsiveness to RNA-LPX vaccination was influenced by the organ-specific microenvironment. This question is particularly relevant because the liver is a major metastatic site in SCLC^32^, and RNA-LPX vaccination could potentially target both primary lung tumors and distant metastases. Importantly, vaccination conferred comparable survival benefits in mice bearing SCLC-A tumors in the lung or in the liver, even when tumors were established by intrapulmonary inoculation. In both anatomical sites, RNA-LPX vaccination increased antigen-specific T cells and induced changes in myeloid populations expressing co-stimulatory molecules such as CD40 and CD86. These findings suggest that the limited responsiveness of SCLC-A is not primarily determined by the anatomical site of engraftment in this model, but may instead reflect tumor-intrinsic mechanisms of immune resistance.

Multiome snRNA-seq and snATAC-seq of SCLC-A tumors in the lung and in the liver revealed enhanced prevalence of signatures associated with cytotoxic T-cell responses following vaccination, consistent with increased regulatory activity involving TFs such as *Eomes* and *Tbx21*, after RNA vaccination consistent with prominent vaccine-elicited cytotoxic T-cell responses. Moreover, differential chromatin accessibility and regulatory analyses revealed extensive transcriptional and epigenetic remodeling in myeloid and non-immune compartments, involving TFs associated with myeloid/APC activation, such as *Cebpb* and interferon signaling (*Stat1*, *Irf1*, *Irf7*). Differential expression analysis showed increased expression of *Cxcl9* in livers of vaccinated mice, which may promote T-cell infiltration and has been linked to responsiveness to checkpoint inhibitors. Together, these changes suggest a broader inflammatory response across multiple cellular compartments that may support T and NK cell-mediated tumor control.

However, despite these vaccination-induced changes in the TME, therapeutic responsiveness of SCLC-A to RNA-LPX vaccination was limited, suggesting that additional immune stimulation might be required to achieve effective tumor control. We therefore investigated the potential of IFN-α together with IL-15-sushi to enhance antigen presentation and innate immune activation in our model.

Cytokines, such as IFN-α and IL-2 have long been investigated as cancer immunotherapies, although their clinical application has been limited by systemic toxicity^33,34^. Local delivery of RNA-encoded cytokines offer a promising alternative^18,19^, especially in approaches that enable more targeted cytokine expression^35,36^. In our study we used IFN-α together with IL-15-sushi, an IL-15–IL-15Rα sushi-domain fusion that promotes NK and CD8^+^ T-cell responses while avoiding some of the limitations of IL-2, particularly regulatory T-cell expansion and activation-induced cell death of effector T cells^37–39^.

Importantly, combining RNA-LPX vaccination with liver-targeted RNA-encoded IFN-α and IL-15-sushi markedly improved survival compared with either treatment alone. Although vaccination alone elicited high frequencies of antigen-specific CD8 T cells, this T-cell response did not translate into comparable therapeutic efficacy, suggesting that effective T-cell priming alone was insufficient to overcome immune resistance in SCLC-A. These findings highlight the importance of addressing both tumor-intrinsic limitations and immunosuppressive mechanisms within the TME.

Mechanistically, combination treatment broadened CD8 T-cell activation beyond Rpl18- specific cells, suggesting a more diversified immune response that may help limit antigen-dependent tumor escape. Cytokine treatment enhanced MHC-I expression by tumor cells, potentially improving antigen presentation and susceptibility to T-cell recognition. In parallel, myeloid populations acquired a more inflammatory phenotype, which may further support antigen presentation and effector-cell activity. These changes may be particularly relevant given the association between suppressive myeloid populations and poor immunotherapy responses^15^. Together, these findings suggest that cytokine delivery complements RNA-LPX vaccination by enhancing tumor immunogenicity and promoting a more permissive microenvironment for antitumor immune responses.

Overall, our work indicates that immune hot SCLC-Y is relatively susceptible to RNA-LPX vaccination across a range of antigen strengths, whereas SCLC-A requires more effective antigen targeting and additional immune modulation to achieve tumor control. RNA-encoded cytokines partially overcome these barriers by inducing tumor-cell MHC-I upregulation, broadening T-cell activation, and reprogramming myeloid cells. These findings support the development of subtype-specific strategies that combine rational antigen selection with cytokine RNA-based immune modulation to overcome immunotherapy resistance in SCLC-A. Importantly, our murine models recapitulate distinct immune phenotypes of human SCLC across different anatomical sites, providing a valuable preclinical platform to investigate tumor– immune interactions and evaluate novel therapeutic strategies with translational potential.

## MATERIALS & METHODS

### Ethical statement

All animal experiments were conducted in accordance with the local authorities of Rhineland-Palatinate.

### Cell culture

SCLC cell lines were isolated from tumor-bearing lungs of fully backcrossed male *Rb1^fl/fl^ Tp53^fl/fl^* C57BL/6 mice. Cell lines were generously donated by Dr. Filippo Beleggia from the Department of Translational Genomics of Colone University Medical Center. Cells were cultured as adherent cells or spheroids in DMEM/F12 (1:1, Gibco, Cat# 11320033) + 2.5 % FBS (Bio & Sell, Cat# FBS. S 0615) + 2 mM L-Glutamin (Gibco, Cat# 25030081) + 30 nM Sodium Selenite (Sigma-Aldrich, Cat# S9133-1MG) + 10 µg/mL Apo-Transferrin (Sigma-Aldrich, Cat# T1147-100MG) + 5 µg/mL human zinc-insulin (Gibco, Cat# 12585014) + 10 nM Hydrocortisone (Sigma-Aldrich, Cat# H0135-1MG) + 10 nM β-Estradiol (Sigma-Aldrich, Cat# E2758-250MG).

### Neoantigen introduction to mSCLC cell lines

Neoantigens were transfected using lentiviral particles. Lentiviral supernatants with a VSV-G envelope were generated in HEK293T-17 cells. Cells were seeded in a density of 7 x 10^5^ cells/cm^2^ in DMEM (Glutamax, Gibco, Cat# 10566016) + 10 % FBS. On the following day, cells were analyzed to ensure confluency as well as equal distribution in the culture dish (145 cm^2^, ThermoScientific, Cat# 150468). All reagents were allowed to accommodate room temperature. Per dish, 7.5 mL of Opti-Mem (Gibco, Cat# 31985070) are mixed with 12 µg of lentiviral transfer vector, 2.3 µg of VSV-G and 16 µg of Gag-Pol (pCMVΔR8.91) vector in a 50 mL reaction tube. 90 µl Transit-LT1 (Mirus, Cat# MIR2034) is added and quickly mixed by pipetting. The transfection mix is incubated for 20 min at room temperature and added to the cells by pipetting. Cells are transfected for 16 to 18 h at 37 °C and 5 % CO_2_. On the following day, medium is changed. On day 4 of the generation, supernatant is collected and filtered through a 0.45 µM sterile filter and collected in a 2 mL cryovial.

In preparation for lentiviral transduction, untreated 24-well plates are coated with 250 µl retronectin (Takara, Cat# T100A) working solution (20 µg/mL in PBS) per well over night at 4 °C. On the following day, retronectin solution is aspirated, wells are washed with PBS and blocked for 30 min. at RT using 2 % BSA (w/v, Sigma-Aldrich, Cat# A1470-25G). Wells are washed again with PBS and 250 µl viral supernatants are added onto the wells. The plate is centrifuged for 15 min at 1500 x g and 15 °C. The supernatant is aspirated and the steps are repeated two more times. After the last spin, supernatant is aspirated and wells are washed again with PBS. Target cells are seeded in a concentration of 40.000 cells/well in 1 mL Medium and are incubated for 2 days at 37 °C, 5 % CO2. Infection efficiency is validated via Flow Cytometry.

### Animal experiments

Animals were kept under specific-pathogen free conditions. Mice were housed in individually ventilated cages at ∼ 75 air changes per hour. Relative temperature was set to 20 – 24 °C, relative humidity was set to 45 – 65 %. Animals were fed ad libitum throughout all experiments. A 12 h:12 h light-dark cycle was implemented. For all experiments 8 – 12 weeks old C57BL/6_OlaHsd (Envigo, Cat# 5703M) or female Nod scid-gamma (NSG, Charles River, Cat# 614) mice were used.

All experiments were performed according to Directive 2010/63/EU, animal care was in accordance with institutional guidelines. Termination criteria for in vivo experiments were established in advance and approved by the local animal regulatory authority. These included, for example, >20% body weight loss, a body conditioning score of 2 or lower, visibly impaired or increased abdominal breathing for lung models, abdominal swelling for liver tumor models and additional criteria related to the animals’ overall condition and behavior as defined by a pre-established scoring system. Survival endpoints were determined according to these predefined criteria, and lung tumor burden was macroscopically evaluated to verify that mice had reached the experimental endpoint. Animals were censored in survival analyses if sacrifice or death occurred for reasons unrelated to lung tumor burden. Symbols shown in survival curves represent both events and censored observations. Mice were randomly allocated to either treatment or control groups. Depending on the tumor model and experimental design, group sizes generally consisted of 10 mice per group for pilot studies, 12 (SCLC-A) or 15 (SCLC-Y) mice per group for monotherapy and 17 (SCLC-A) or 22 (SCLC-Y) mice for combination therapy for survival studies and 6 (SCLC-A) or 8 (SCLC-Y) mice per group for flow cytometric analyses, as specified in the respective figure legends. Sample size calculations were typically performed using G*Power based on previously established biologically meaningful effect sizes and were approved by the local regulatory authority.

### Tumor models and treatments

For tumor inoculation, mice were either injected with 4 - 5 x 10^6^ SCLC tumor cells in 125 µl PBS (Hi-Media, Cat# TL1006) via tail vein injection or with 2 x 10^6^ cells intrapulmonal in 30 µl 1:1 PBS:Matrigel (Corning, Cat# 354234). Intrapulmonary Injections were performed into the right thorax under ongoing isofluorane anesthesia. Mice were randomly assigned to therapy groups and monitored 3 x per week after tumor inoculation. RNA-LPX or formulated RNA-encoded cytokines were intravenously injected in 100-200µl. All treatments were conducted according to the respective experimental layout.

For analysis of circulating immune cells, up to 60 µl blood was isolated via facial vein puncture and collected in heparin-coated tubes (Sarstedt, Cat# 20.1292) until further processing. For analysis of liver enzymes in blood, 150 µl of blood was withdrawn retro-orbitally while mice were under 2.5 % isoflurane anesthesia. Blood was collected in serum tubes (Sarstedt, Cat# 20.1343.100) until further processing.

### Mutation selection and prioritization

Whole exome DNA-sequencing reads were aligned to the mm9 genome using bwa^40^ (default options, v0.7.10). Bulk RNA-seq reads were aligned to the mm9 reference using STAR^41^ (default options, v2.1.4a).

Strelka2^42^; default options for whole exome sequencing, v2.0.14) was used to call somatic SNVs and short insertion/deletion (InDels) on each cell line. Only passed calls were considered for downstream analyses.

Transcript abundance estimation was performed with sailfish quant^43^ (default options, v0.7.6). The resulting transcripts per million (TPM) values were converted into RPKM values:

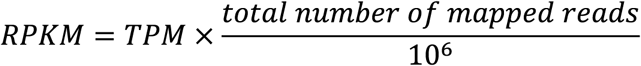

Non-synonymous SNVs and InDels were investigated in the context of the respective 27mer amino acid neoantigen candidate with the mutated amino acid in the center (position 14). Frameshift candidates were extended until reaching the next stop codon.

Neoantigen candidates were subsequently annotated with NeoFox^44^ (v0.6.1) to analyse immunogenicity-related features such as Vaxrank^24^.

### RNA

Sequences encoding the cytokines (murine IL-15 receptor-α sushi domain linked to murine IL-15 or murine IFN-α4), were cloned into plasmid vectors harboring a 5’ untranslated region (UTR), a 3’ UTR and poly A tail ^45–47^. Following linearization of the DNA template, RNA was synthesized *in vitro* transcription (IVT) with T7 RNA polymerase (Thermo Fisher Scientific) in the presence of adenosine 5′-triphosphate, cytidine 5′-triphosphate, guanosine 5′-triphosphate and 1-methylpseudouridine-5’-triphosphate (m1ΨTP; Thermo Fisher Scientific) substituting for uridine-5’-triphosphate (UTP) as previously described^48,49^. CleanCap N-7413 (TriLink Biotechnologies) was introduced co-transcriptionally. To remove double-stranded RNA contaminants, cellulose purification was performed as previously described^49^.

Predicted tumor antigen candidates and antigens/Rpl18 were cloned into plasmid templates for in vitro transcription, up to 10 antigens separated by GS linkers and followed by a poly-A tail.

### RNA formulation

RNA-LPX for antigen expression were generated as previously described^31^.

RNA-encoded cytokines (max. 3 µg RNA in 200 µl) were formulated in TransIT (Mirus, Cat# MIR2250) according to the manufacturers protocol and injected via tail vein.

### Enzyme-analysis via INDIKO

Liver damage was assessed by analysis of systemic liver damage associated enzymes. Serum was collected by centrifuging blood collected in serum tubes for 5 min at 10.000 *x g* to separate serum. Serum is diluted 1:1 using water and serum levels are measured by spectroscopic measurements using Indiko^TM^ chemical analyzer

### Single cell isolation from organs

For subsequent analysis in flow cytometry and single-cell sequencing, single cells were isolated from organs. Isolation of splenocytes was performed by enzymatic digestion. Spleens were minced using scissors and topped with 2.5 mL digestion buffer (1 mg/mL Collagenase D [Sigma-Aldrich, Cat# 11088858001) + 50 µg/mL DNAse I [Sigma-Aldrich, Cat# 1128493200] in PBS). Spleens were further prepared using gentleMACS Octo-Dissociator according to the manufacturers protocol. Cell suspension is filtered through a 70 µM cell strainer (Greiner Bio- One, Cat# 542070) and toppled with RBC lysis buffer (150 mM NH_4_Cl, 10 mM KHCO_3_, 0.1 mM EDTA, pH 7.2 – 7.4) for 5 min to lyse erythrocytes. Cells are washed and resuspended in PBS + 5 % FCS + 50 mM EDTA (Sigma-Aldrich, Cat# 03690-100 mL) until further processing. Cells were isolated from lungs by using gentleMACS Octo-Dissociator according to the manufacturers protocol. Before running the program, lungs were minced manually into smaller pieces using scissors. After generating a single cell suspension, lysis of red blood cells and washing was performed according to the protocol for splenocyte isolation.

Before isolating single cells from liver, livers were perfused using up to 40 mL perfusion buffer (HBSS [Gibco, Cat# 14025092] + 2 mM EDTA) via portal vein after euthanasia. Gallbladder is removed before isolating single cells. Perfused livers were dissolved using gentleMACS according to the manufacturers protocol. Isolated cells were separated using a 70 µM cell strainer and were subsequently toppled once with 5 mL DMEM without serum (Hi-Media, Cat# AL007S), centrifuged once for 10 min at 300 x g and washed afterwards with 40 mL PBS. Cell suspension is split equally for further processing. An isotone percoll working solution was prepared using a 1:10 dilution of 10 x PBS (Gibco, Cat# 70011044) and percoll (Cytiva, Cat# 17-0891-01). Isotone percoll is mixed with DMEM in order to prepare a density gradient. 3 mL high percentage percoll is prepared in a 15 mL reaction tube. Cells were resuspended in 5 mL low percentage percoll and carefully layered on the high percentage percoll. For isolation of lymphoid cells, cell suspension is passed through a 40:80 gradient, for isolation of myeloid compartment cells cell suspension is passed through a 25:50 gradient by centrifuging at 1350 x g for 30 min without breaking. The interphase is collected and cells are washed with PBS + 5 % FCS + 2 mM EDTA before further processing.

### Synthetic peptides

For ex vivo re-stimulation during immunogenicity testing, peptides containing non-synonymous mutations were synthesized as 27-mer peptides, containing the mutated amino acid at center position or wildtype control peptides. Biosynth synthesized all peptides with a purity above 70%.

### Enzyme-linked immunospot (ELISPOT) assay for immunogenicity testing of tumorantigen candidates

ELISpot was performed assessing T-cell reactivity by measuring IFN-γ secretion. 96-well microfilter plates (Merck, MSIPS4510) were activated with 35 % EtOH (Carl Roth, Cat# P075.1), washed 3 x with 150 µl PBS/well and coated with 10 µg/ml anti-mouse IFN-γ (Mabtech, Cat# 3321-3-250) over night at 4 °C. Plates were washed 3 x with 150 µl PBS/well and were blocked for 1 – 4 h using RPMI + Glutamax + 10 % FBS + 1 mM Sodium Pyruvate + 1 x MEM NEAA + 0.5 % Pen-Strep + 50 µM β-Mercaptoethanol. 5 x 10^4^ cells/well of target cells or BMDCs were seeded together with either 1 x 10^5^ CD4^+^ T cells, 5 x 10^4^ CD8^+^ T cells or 5 x 10^5^ full splenocytes. In case of testing against a peptide, BMDCs were loaded with 5 µg/ml per peptide. As positive control, effectors were incubated with 2 µg/ml ConA (Invitrogen, Cat# 00-4978-93). Cells were incubated for 16 – 24 h at 37 °C/5 % CO_2_. Cells were discarded, plates were washed 3 x with 150 µl PBS/well and per well 2 µg/ml IFN-γ secondary antibody (Mabtech, Cat# 3321-6-250) in 0.5 % BSA/PBS (Sigma-Aldrich, Cat# A1470-25G) were added for 2 h at 37 °C. Wells were washed as previously described and treated for 1 h with 1 µl/mL ExtrAvidin-Phosphatase (Sigma-Aldrich, Cat# E2636-.2ML). Wells were washed as previously described and 100 µl of BCIP/NBT chromogenic solution (Sigma-Aldrich, Cat# B1911-100ML) was added per well for 7 minutes. Enzymatic reaction was stopped by flushing wells with water. Wells were captured and spot counts were analyzed using CTL Immunospot S6 Ultimate reader.

### Histology

Tumor tissue were harvested from mice, fixed with 4 % histofix overnight at 4 °C and embedded in paraffin blocks (FFPE). Cell pellets were collected by centrifugation at 300 × g for 5 min, fixed in 4 % histofix for 1 h at RT, embedded in HistoGel and subsequently paraffin- embedded.

For chromogenic IHC, tissue sections were stained with anti-ASCL1 (Abcam, ab211327, 1:1000), anti-YAP (CST, 17074, 1:200), and anti-GFP (Abcam, ab183734, 1:500) using the Leica BOND Rx autostainer. The standard IHC protocol adjusted to 30 min primary antibody incubation with HIER: 20min ER2 was used. The staining was detected using BOND Polymer Refined Detection Kit (Leica, DS9800). Images were acquired using Aperio AT2 Slide Scanner (Leica Biosystems).

### Phenocycler Fusion assay

The PhenoCycler-Fusion system (Quanterix, formerly Akoya Biosciences) was used to characterize the SCLC tumor microenvironment. Sample preparation was performed according to the PhenoCycler-Fusion user manual (PhenoImager Fusion SW Version: 2.1.0). Briefly, 3 μm FFPE tissue sections were pre-processed by deparaffinization, dewaxing/rehydration, and heat-mediated antigen retrieval (Akoya Biosciences, 7000017). Successive sample hydration, antibody staining and post-staining fixation were performed using the Parhelia Spatial Station. Tissues were simultaneously stained with the barcoded-antibody panel (see **Antibody list for Histology**) followed by signal stabilization (post-staining fixation to preserve barcode-epitope complexes). Cyclic reporter hybridization (fluorescent oligonucleotide binding to antibody barcodes) and following high-plex multiplexed images were acquired on the PhenoImager Fusion platform using Fusion Software.

### Antibody list for Histology

| **Antibody** | **Cat.No.** | **Dilution** | **Fluorophore** | **Cat.No.** | **Dilution** |
|---|---|---|---|---|---|
| F4/80-BX006 | Akoya Biosciences, 200082 | 1:200 | RX006-AF647 | Akoya Biosciences, 6550026 | 1:50 |
| CD8-BX026 | Akoya Biosciences, 200092 | 1:200 | RX026-ATTO550 | Akoya Biosciences, 6250007 | 1:50 |
| Ki67-BX047 | Akoya Biosciences, 232179 | 1:200 | RX047-ATTO550 | Akoya Biosciences, 6250012 | 1:50 |

### Image analysis

Image analysis was performed using QuPath (Version 0.6.0)^50^. Tissues were segmented into tumor and stromal regions based on a thresholder for Ki67 and manual correction. TLS-like structures were excluded from analysis. Cells were successively segmented using the StarDist plugin (Schmidt, U., Weigert, M., Broaddus, C., Myers, G., 2018). For each marker, object classifiers were independently developed using a representative training image compiled from all samples to avoid bias. Classifiers were based on characteristic features such as signal intensity patterns, subcellular localization, and morphological criteria. Trained classifiers were applied sequentially to each segmented sample. Data were exported from QuPath and processed using R studio (Version 2024.04.0) and Microsoft Excel (Microsoft 365 MSO, Version 2507). Representative images were generated using the ImageJ plugin of QuPath with median filter (radius 1).

### Flow cytometry

In case of staining blood, 50 µl were added to a well of a 96-well v-bottom plate. 100 µl RBC lysis buffer was added for 1.5 min and cells were centrifuged for 3.5 minutes at 380 x g, 4 °C. Cells were washed once with flow buffer (PBS + 5 % FBS + 2 mM EDTA), centrifuged and washed once with PBS before continuing staining.

Cell suspensions were generated as described above, were washed with flow buffer and transferred into a 96-well v-bottom plate (Corning, Cat# 3894) for staining. 1.5 x 10^6^ cells were inserted per staining. Cells were stained with 50 µl Tetramer-solution (MBLBio, Cat# MBL-TC-5516-2) for 20 min at 2 – 8 °C and then topped with 50 µl of 2 x concentrated extracellular mastermix for further 20 min at 2 - 8 °C. Cells were washed once with PBS before addition of 100 µl diluted viability dye and, if applicable, streptavidin-coupled dye. Solution was incubated for 15 min at 2 – 8 °C before centrifuging and washings cells twice with PBS. Cells were fixed and permeabilized by adding 200 µl of Fix/Perm solution (Invitrogen, Cat# 000-5523-00), prepared according to the manufacturers instructions, onto the cells. Cells are incubated with the solution for 30 min at 2 – 8 °C. Meanwhile, permeabilization buffer (Perm-buffer) was prepared according to the manufacturers instruction and antibodies for intracellular/intranuclear staining were diluted. Cells were washed once in Perm-buffer, 100 µl of antibody solution was added and incubated for further 30 minutes. Afterwards cells were washed twice with Perm-buffer and taken up in 1:10 diluted counting beads in flow buffer before acquisition.

### Antibody list for flow cytometry

| Antibody | Citation |
|---|---|
| APC anti-mouse H-2Db | BioLegend Cat# 111514, RRID:AB_2565862 |
| APC/Cyanine7 anti-mouse CD11c | BioLegend Cat# 117324, RRID:AB_830649 |
| APC/Cyanine7 anti-mouse/rat CD29 | BioLegend Cat# 102225, RRID:AB_2128078 |
| Biotin anti-mouse CD127 (IL-7Rα) | BioLegend Cat# 135005, RRID:AB_1953262 |
| Biotin anti-mouse CD19 | BioLegend Cat# 115503, RRID:AB_313638 |
| Biotin anti-mouse CD3 | BioLegend Cat# 100243, RRID:AB_2563946 |
| Biotin anti-mouse CD49b (pan-NK cells) | BioLegend Cat# 108903, RRID:AB_313410 |
| Biotin anti-mouse CD5 | BioLegend Cat# 100603, RRID:AB_312732 |
| Biotin anti-mouse CD64 (FcγRI) | BioLegend Cat# 139318, RRID:AB_2566557 |
| Biotin anti-mouse F4/80 | BioLegend Cat# 123106, RRID:AB_893501 |
| Biotin anti-mouse FcεRIα | BioLegend Cat# 134303, RRID:AB_1626100 |
| Biotin anti-mouse Ly-6G | BioLegend Cat# 127604, RRID:AB_1186108 |
| Biotin anti-mouse TER-119/Erythroid Cells | BioLegend Cat# 116203, RRID:AB_313704 |
| Brilliant Violet 421(TM) anti-mouse CD192 (CCR2) | BioLegend Cat# 150605, RRID:AB_2571913 |
| Brilliant Violet 421(TM) anti-mouse CD25 | BioLegend Cat# 102034, RRID:AB_11203373 |
| Brilliant Violet 570(TM) anti-mouse CD62L | BioLegend Cat# 104433, RRID:AB_10900262 |
| Brilliant Violet 570(TM) anti-mouse Ly-6C | BioLegend Cat# 128029, RRID:AB_10896061 |
| Brilliant Violet 605(TM) anti-mouse Ki-67 | BioLegend Cat# 652413, RRID:AB_2562664 |
| Brilliant Violet 650(TM) anti-human CD4 | BioLegend Cat# 317436, RRID:AB_2563050 |
| Brilliant Violet 650(TM) anti-mouse NK-1.1 | BioLegend Cat# 108736, RRID:AB_2563159 |
| Brilliant Violet 650(TM) anti-mouse/rat XCR1 | BioLegend Cat# 148220, RRID:AB_2566410 |
| Brilliant Violet 711(TM) anti-T-bet | BioLegend Cat# 644819, RRID:AB_11218985 |
| Brilliant Violet 750(TM) anti-mouse/human CD44 | BioLegend Cat# 103079, RRID:AB_2941373 |
| CD101 Monoclonal Antibody (Moushi101), PE-Cyanine7, eBioscience | (Thermo Fisher Scientific Cat# 25-1011-80, RRID:AB_2573377) |
| CD103 (Integrin αIEL) | BD Biosciences Cat# 564322, RRID:AB_2738744 |
| CD103 (Integrin αIEL) | BD Biosciences Cat# 565529, RRID:AB_2739282 |
| CD119 (IFN-γ Receptor α) | BD Biosciences Cat# 740032, RRID:AB_2739804 |
| CD11b Monoclonal Antibody (M1/70), Brilliant Ultra Violet™ 805, eBioscience | (Thermo Fisher Scientific Cat# 368-0112-80, RRID:AB_2896081) |
| CD19 | BD Biosciences Cat# 563333, RRID:AB_2738141 |
| CD27 | BD Biosciences Cat# 746742, RRID:AB_2744005 |
| CD274 (PD-L1, B7-H1) Monoclonal Antibody (MIH5), PE-Cyanine7, eBioscience | (Thermo Fisher Scientific Cat# 25-5982-82, RRID:AB_2573509) |
| CD317 (BST2) | BD Biosciences Cat# 747600, RRID:AB_2744168 |
| CD366 (TIM-3) | BD Biosciences Cat# 747620, RRID:AB_2744186 |
| CD40 Monoclonal Antibody (HM40-3), eFluor™ 450, eBioscience | (Thermo Fisher Scientific Cat# 48-0402-82, RRID:AB_2574017) |
| EOMES Monoclonal Antibody (Dan11mag), PE-Cyanine5, eBioscience | (Thermo Fisher Scientific Cat# 15-4875-80, RRID:AB_2802206) |
| F4/80 Monoclonal Antibody (BM8), APC, eBioscience | (Thermo Fisher Scientific Cat# 17-4801-82, RRID:AB_2784648) |
| FITC anti-mouse CD206 (MMR) | BioLegend Cat# 141704, RRID:AB_10901166 |
| FOXP3 Monoclonal Antibody (FJK-16s), eFluor™ 450, eBioscience | (Thermo Fisher Scientific Cat# 48-5773-80, RRID:AB_1518813) |
| I-A/I-E | BD Biosciences Cat# 743871, RRID:AB_2741822 |
| KLRG1 Monoclonal Antibody (2F1), APC-eFluor™ 780, eBioscience | (Thermo Fisher Scientific Cat# 47-5893-82, RRID:AB_2573988) |
| Ly-108 Monoclonal Antibody (eBio13G3-19D (13G3-19D)), NovaFluor™ Yellow 700, eBioscience | Thermo Fisher Scientific Cat# M050T02Y06, RRID:AB_2931047 |
| Ly-6G Monoclonal Antibody (1A8-Ly6g), PerCP-eFluor™ 710, eBioscience | (Thermo Fisher Scientific Cat# 46-9668-82, RRID:AB_2573893) |
| Mouse H60 PE-conjugated Antibody | R and D Systems Cat# FAB1155P, RRID:AB_2115298 |
| Mouse MULT-1 Phycoerythrin MAb (Clone 237104) | (R and D Systems Cat# FAB2588P, RRID:AB_10973344) |
| Mouse Rae-1 Pan Specific PE-conjugated Antibody | R and D Systems Cat# FAB17582P, RRID:AB_357086 |
| Mouse TCF7/TCF1 Alexa Fluor® 488-conjugated Antibody | R and D Systems Cat# IC8224G, RRID:AB_3656680 |
| Ms CD115 (CSF-1R) BUV661 AFS98 50ug | BD Biosciences Cat# 750949, RRID:AB_2875030 |
| Ms CD172a BUV563 P84 50ug | BD Biosciences Cat# 741349, RRID:AB_2870850 |
| Ms CD274 BUV615 MIH5 50ug | BD Biosciences Cat# 752339, RRID:AB_2875856 |
| Ms CD279 (PD-1) BV786 RMP1-30 50ug | BD Biosciences Cat# 748264, RRID:AB_2872692 |
| Ms CD3 MolCpx BUV563 17A2 50ug | BD Biosciences Cat# 741319, RRID:AB_2870837 |
| Ms CD31 BV750 390 50ug | BD Biosciences Cat# 746871, RRID:AB_2871671 |
| Ms CD44 BUV563 IM7 50ug | BD Biosciences Cat# 569705, RRID:AB_3626317 |
| Ms CD45 BUV496 30-F11 50ug | BD Biosciences Cat# 569673, RRID:AB_3675211 |
| Ms CD69 BUV615 H1.2F3 50ug | BD Biosciences Cat# 751593, RRID:AB_2875587 |
| Ms CD86 RB780 PO3 50ug | BD Biosciences Cat# 755499, RRID:AB_3687863 |
| Ms CD8a RB705 5H10-1 50ug | BD Biosciences Cat# 757841, RRID:AB_3689997 |
| Ms H-2Kb BUV563 AF6-88.5 50ug | BD Biosciences Cat# 748823, RRID:AB_2873226 |
| Ms H-2Kb BV421 AF6-88.5 25ug | BD Biosciences Cat# 566240, RRID:AB_3683733 |
| Ms KLRG1 BUV661 2F1 50ug | BD Biosciences Cat# 741586, RRID:AB_2870999 |
| Ms NK-1.1 BUV737 PK136 50ug | BD Biosciences Cat# 741715, RRID:AB_2871088 |
| PE anti-mouse CD64 (FcγRI) | BioLegend Cat# 139304, RRID:AB_10612740 |
| PE anti-mouse IFN-γ R β chain | BioLegend Cat# 113603, RRID:AB_313560 |
| PE/Cyanine5 anti-mouse CD69 | BioLegend Cat# 104509, RRID:AB_313112 |
| PE/Cyanine7 anti-mouse CD366 (Tim-3) | BioLegend Cat# 119715, RRID:AB_2571932 |
| PE/Dazzle(TM) 594 anti-mouse CD170 (Siglec-F) | BioLegend Cat# 155529, RRID:AB_2890716 |
| PE/Dazzle(TM) 594 anti-mouse CD69 | BioLegend Cat# 104536, RRID:AB_2565583 |
| PE/Dazzle™ 594 anti-mouse CD127 (IL-7Rα) | BioLegend Cat# 135032, RRID:AB_2564217 |
| PE/Fire(TM) 810 anti-mouse CX3CR1 | BioLegend Cat# 149057, RRID:AB_2936671 |
| Rat Anti-CD25 Monoclonal Antibody, Phycoerythrin Conjugated, Clone PC61 | BD Biosciences Cat# 553866, RRID:AB_395101 |
| Rat Anti-CD90.2 Monoclonal Antibody, FITC Conjugated, Clone 53-2.1 | BD Biosciences Cat# 553003, RRID:AB_394542 |
| Rat/Ms CD49a BUV805 Ha31/8 50ug | BD Biosciences Cat# 741976, RRID:AB_2871280 |
| Spark Red™ 718 anti-mouse CD192 (CCR2) (Flexi-Fluor™) | (BioLegend Cat# 285110, RRID:AB_3106362) |
| Spark UV(TM) 387 anti-mouse I-A/I-E | BioLegend Cat# 107669, RRID:AB_2922456 |
| T-bet RB780 O4-46 25Tst | BD Biosciences Cat# 569090, RRID:AB_3684770 |
| TIM-4 | BD Biosciences Cat# 742777, RRID:AB_2741041 |
| TOX Antibody, anti-human/mouse, PE, REAfinity™ | Miltenyi Biotec Cat# 130-120-716, RRID:AB_2801780 |
| VISTA Monoclonal Antibody (MIH64), Brilliant Ultra Violet™ 737, eBioscience | (Thermo Fisher Scientific Cat# 367-1083-80, RRID:AB_2895988) |

### Streptavidin for flow cytometry

| **Streptavidin** | **Citation** |
|---|---|
| SparkBlue™ 574 Streptavidin | Biolegend Cat# 405355 |
| BD Horizon™ BV510 Streptavidin | BD Biosciences Cat# 563261 |

### Viability dyes for flow cytometry

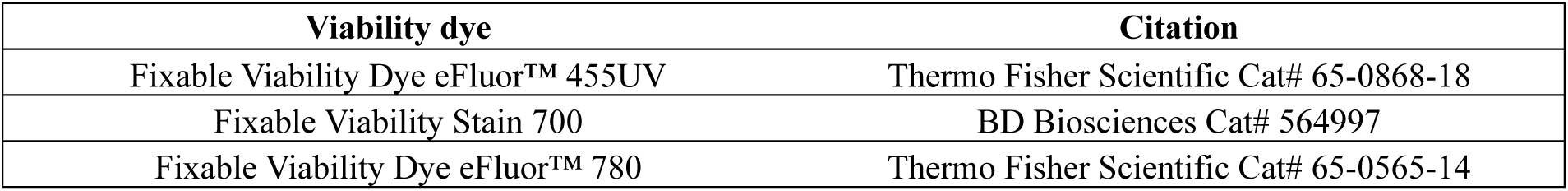

### In vitro IFN-γ treatment

Spheroid cell lines were separated by incubating cells for 10 min in Accutase (Bio & Sell, Cat# BS.L2193). Single cells were seeded in a density of 3 x 10^4^ cells per well. For adherent cells, cells were seeded in a density of 3 x 10^4^ cells/cm^2^ for overnight stimulation or 1 x 10^4^ cells/cm^2^ for prolonged stimulation. Cells were treated with 50 ng/mL murine IFN-γ (Fisher Scientific, Cat# 17872833) by adding directly to the medium.

### DNA & RNA extraction from cells & fresh frozen tissue

According to the manufacturers protocol, cells were extracted using Qiagens DNeasy Blood & Tissue Kit for genomic DNA or Qiagens RNeasy Mini Kit for total RNA. Quality and Quantity was assessed using a Qubit fluorometer, the Nanodrop2000 and the Fragment Analyzer 5200/5300.

### NGS library preparation

Whole exome libraries were prepared in replicates with an input of 100 ng genomic DNA (gDNA) or 50 ng total RNA. The gDNA was fragmented to a size range of 200 bp ± 50 bp using the Covaris Ultrasonificator ME220 while the total RNA went through enzymatic fragmentation – the RNA samples had to be primed and reverse-transcripted after fragmentation first. Afterwards all samples were end-repaired and adenylated, followed by the ligation of a full-length single index eight-nucleotide NEXTFLEX DNA barcode (Perkin Elmer). The library was then pre-amplified using the KAPA Hyper Prep kit (Roche) performing 9 cycles for DNA libraries or 14 cycles for RNA libraries. Subsequently, target regions were hybridized to biotinylated baits (SureSelectXT Human All Exon v6 and SureSelect XT Reagent kit, Agilent) and isolated using streptavidin-coated magnetic beads (Invitrogen). The post-capture libraries were then amplified with the KAPA Library Amplification kit (Roche) performing 10 PCR cycles.

All intermediate and final library purification steps for both library types were carried out using AMPure XP beads (Beckman Coulter).

All QC steps, for library preparation, were performed using a Qubit fluorometer (Invitrogen) for quantitative analysis and the 2100 Bioanalyzer Instrument or the Fragment Analyzer 5200/5300 for qualitative analysis.

### (bulk) Sequencing

All libraries were sequenced in paired-end mode (2 x 50 nt) on an Illumina NovaSeq 6000 instrument resulting in around 150 million distinct sequencing reads per whole-exome DNA library or 100 million distinct sequencing reads per whole-exome RNA library.

### Nuclei isolation

For single-nucleus multiome sequencing, nuclei were isolated from freshly isolated cell suspensions using the Chromium Nuclei Isolation protocol (10x Genomics, CG000366 Rev. D). Prior to nuclei isolation, cell viability was assessed using automated cell counting (LUNA- FX7, Logos Biosystems) and brightfield microscopy. Samples displaying reduced viability were subjected to dead-cell removal (CG000093 Rev. C) prior to nuclei isolation according to the manufacturer’s instructions. For lung samples, approximately 8 × 10^5 viable cells were used as input for nuclei isolation. Cells were lysed in 1× lysis buffer for 5 min on ice. For liver samples, approximately 5 × 10^5 viable cells were used as input and lysed in 0.5× lysis buffer for 4 min on ice. Following isolation, nuclei were stained with acridine orange/propidium iodide (AO/PI), quantified using a LUNA-FX7 cell counter, and morphologically assessed by light microscopy. Nuclei suspensions containing debris or aggregates were filtered through 40 μm Flowmi strainers (Carl Roth).

### Single-cell Multiome ATAC + Gene Expression library preparation and sequencing

Single-nucleus ATAC and gene expression libraries were generated using the Chromium Single Cell Multiome ATAC + Gene Expression v1.0 platform (10x Genomics) according to the manufacturer’s instructions. Nuclei were adjusted to the recommended concentration and loaded onto Chromium Next GEM chips targeting recovery of approximately 5,000 nuclei per sample. Following GEM generation, barcoding, transposition, reverse transcription and library amplification, separate ATAC and gene expression libraries were prepared according to the manufacturer’s protocol

Library quality and fragment size distribution were assessed using the Agilent TapeStation system (Agilent Technologies), and library concentrations were measured using the Qubit dsDNA High Sensitivity Assay (Thermo Fisher Scientific). ATAC libraries were pooled based on qPCR-derived molarity estimates obtained using the KAPA Library Quantification Kit (Roche). Libraries were then sequenced on a NovaSeq X instrument (Illumina).

### Single-cell multiome data processing and analysis

Raw sequencing data were processed using Cell Ranger ARC v2.1.0 (10x Genomics), employing the mm10-2020-A-2.0.0 reference genome. Cell Ranger ARC was used for read alignment, barcode assignment, gene expression quantification, chromatin accessibility peak calling, and generation of paired single-nucleus RNA and ATAC count matrices. Sequencing and library quality metrics were assessed using MultiQC.

Downstream analyses were performed in R v4.3.1 using the Seurat (v5.5.1) and Signac (v1.17.1) packages. Low-quality nuclei were removed based on RNA and ATAC quality control metrics, including gene counts, fragment counts, TSS enrichment scores, and nucleosome signal profiles. Gene expression and chromatin accessibility modalities were processed separately and batch-corrected using Harmony (v2.0.5). The Harmony-corrected embeddings were subsequently integrated using Seurat’s weighted nearest neighbor (WNN) framework to obtain a joint multimodal representation. Uniform Manifold Approximation and Projection (UMAP) was used for dimensionality reduction and visualization, and unsupervised clustering was performed for cell population identification. Cluster annotation was based on established marker genes and major cellular compartments, including T/NK cells, myeloid cells, and non-immune populations, were further resolved by iterative subclustering analyses to achieve higher-resolution cell type annotation.

Differential gene expression (DEG) and differential accessibility region (DAR) analyses were conducted to identify condition-associated transcriptional and chromatin changes. Functional enrichment and pathway analyses were performed using clusterProfiler (v4.10.1) and SeuratExtend (v1.2.10). RNA–ATAC concordance was defined as genes that were significantly differentially expressed and had at least one significantly differentially accessible linked region within the same cell compartment. Concordance between transcriptional and chromatin accessibility changes was assessed through integrated multiomic analyses. SCENIC+ (v1.0a2) analysis was performed in Python (v3.11.14) separately for the liver and lung datasets using default parameters. scATAC-seq data was prepared with pycistarget (v2.0a0). For motif enrichment analysis the databases mm10_screen_v10_clust.regions_vs_motifs.rankings, mm10_screen_v10_clust.regions_vs_motifs.scores and motifs-v10nr_clust-nr.mgi-m0.001-o0.0 were used. The resulting eRegulons were filtered to only retain positive regulons (+/+). Extended regulons are derived from indirect motif-to-TF annotation (orthology or motif similarity) and were only kept if no direct regulon was available.

### Quantitative Real-Time PCR using the Fluidigm BioMark HD System

#### Total RNA extraction from mouse tissues

Total RNA was extracted from mouse liver and lung tissues using RNeasy Mini Kit (QIAGEN) following the lipid tissue protocol. Up to 100 mg fresh frozen (FF) tissue were lysed and homogenized in 1 mL Trizol (QIAGEN) using Precellys Evolution Touch Homogenizer (bertin technologies) and lysing kits “Tissue homogenizing CKMix – 2 mL” (bertin technologies CAT. NO.: P000918-LYSK0-A). After homogenization, 200 µL chloroform were added and clean-up was performed according to the manufacturer’s instructions, including on-column DNase digestion (QIAGEN). The total RNA’s concentration, purity and integrity was analyzed using NanoDrop 2000c (Thermo Fisher) and Fragment Analyzer capillary gel electrophoresis instrument (Agilent Advanced Analytical).

#### Reverse Transcription

Reverse transcription was performed using 1 µg total RNA per sample and the PrimeScript™ RT Reagent Kit with gDNA Eraser (TAKARA, Art.No. RR047A) according to the manufacturer’s manual, including gDNA Eraser step. After finalized reverse transcription the cDNA was diluted 1:3 with PCR grade water.

No amplification controls (NAC) of a representative sample collection (n=9) was created following the reverse transcription protocol, replacing the Reverse Transcriptase enzyme with water.

#### Primer Design and Establishment

Primers were designed to address mouse specific mRNA transcripts, by including exon boundaries where possible to prevent genomic binding. Furthermore, the primers were checked for specificity using Primer Blast (NIH – National Library of Medicine - https://www.ncbi.nlm.nih.gov/tools/primer-blast/index.cgi) and UCSC in-silico PCR (University of California Santa Cruz, Genomics Institute -https://genome.ucsc.edu/cgi-bin/hgPcr). To control for potential inhibitory secondary structures within the amplicons, mfold analysis was performed (The UNAfold Web Server -https://www.unafold.org/mfold/applications/dna-folding-form.php). To prevent dimerization the primers were checked using OLIGO ANALYSE TOOL (Eurofins genomics - https://eurofinsgenomics.eu/de/ecom/tools/oligo-analysis/) and OligoAnalyzer (IDT – Integrated DNA Technologies -https://eu.idtdna.com/calc/analyzer).

Primers were established to run at 60°C annealing temperature by individual Quantitative Real-Time PCR (qRT-PCR) including melting curve analysis (Thermo Fisher QuantStudio 3) and were checked for correct and unique PCR products via capillary gel electrophoresis (QIAxcel Advanced – QIAGEN).

Primer efficiencies were controlled by using a dilution series of a suitable control cDNA (preferably Universal Mouse Reference RNA - Ambion (lot# 0006043426) or samples from the experiment) and were accepted with efficiencies of ≥80%.

#### Fluidigm quantitative Real-Time PCR

qRT-PCR was performed using the BioMark HD system (Fluidigm) with SsoFast EvaGreen Supermix with Low ROX (Bio-Rad Laboratories). Samples and assays were prepared and analyzed according to the “Fast Gene Expression Analysis Using EvaGreen on the BioMark or BioMark HD System” Advanced Development Protocol 37. A 96.96 Gene Expression Dynamic Array IFC was loaded using the IFC Controller HX. The thermal protocol followed the manufacturer’s setup and used an annealing temperature of 60°C.

Samples run in dilution series on the Gene Expression Dynamic Array IFC were used to calculate primer efficiencies which were used for the analyses. Each sample was run in technical duplicates, furthermore all assays were run in technical duplicates, resulting in quadruplicates per sample/assay combination. Target gene expression is given in arbitrary expression units and was calculated using Δ Δ Ct-method normalizing to the reference genes: Hprt (Hypoxanthine phosphoribosyltransferase [mus musculus]), Tbp (TATA box binding protein [mus musculus]), Rpl13a (ribosomal protein L13A [mus musculus]), Hmbs (hydroxymethylbilane synthase [mus musculus]) and Ppib (peptidylprolyl isomerase B [mus musculus]).

### Primer table

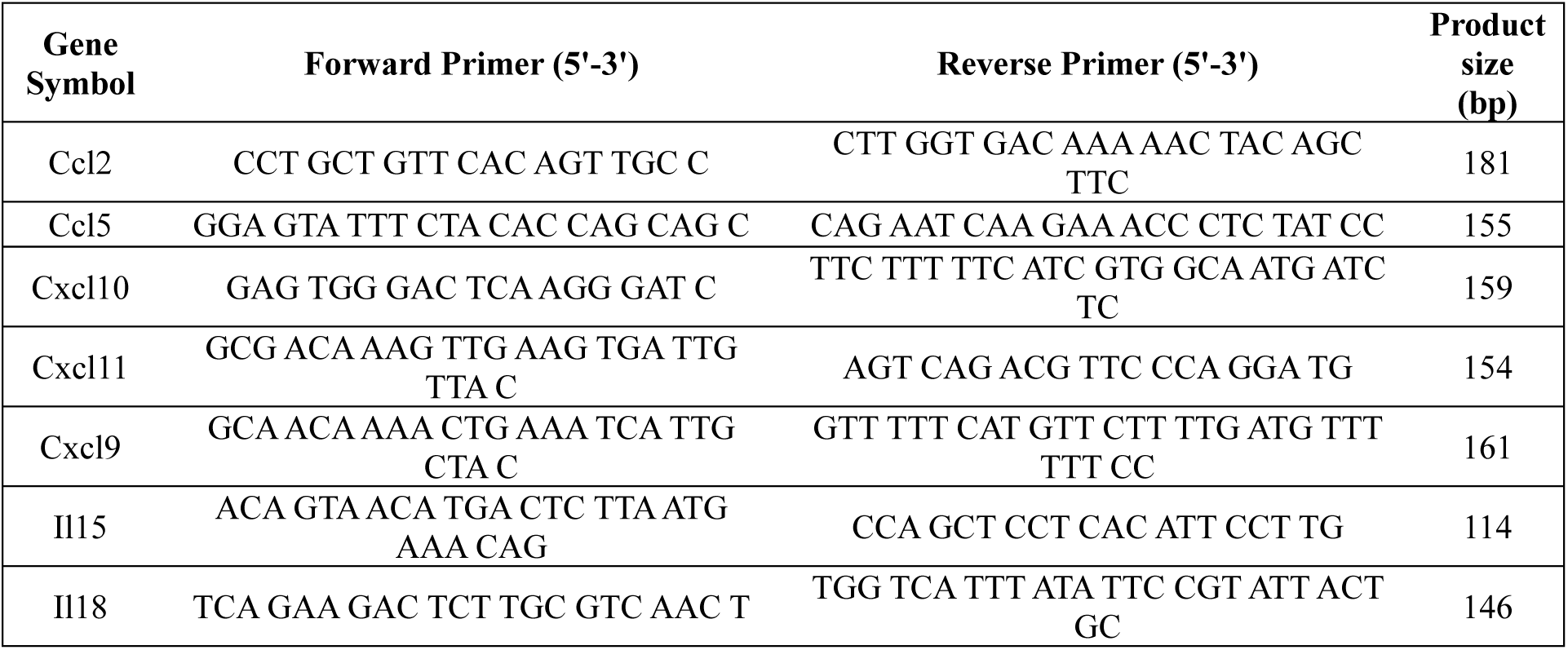

### Multidimensional flow analysis

For high-dimensional flow cytometric analysis of lymphoid immune subsets, liver single-cell suspensions were generated and stained as described above. Samples were acquired on a FACSymphony A5 SE analyzer (Waters Biosciences) equipped with five lasers and fcs.files were imported in FlowJo version 10.10. The files were manually processed to eliminate irrelevant events (debris, doublets and dead cells) and select CD45^+^lineage^-^cells. Following appropriate biexponential transformation of the axes, fcs files were annotated according to the experimental treatment group. Approximately 6000 events of total CD3+ and NK1.1+ cells from each fcs.file were selected and concatenated in a single fcs. file. For the unsupervised identification of lymphoid clusters, the XShift^51^ algorithm was used to facilitate the discovery of rare cell populations with the number of κ-nearest neighbors set to 29. The quality of the generated clusters was subsequently evaluated using the Euclid plugin, comprising two algorithms that asses the presence of potential additional subpopulations (ClusterRCheck /under-clustering) and relatedness/similarity of the identified populations (Taylor Index/over- clustering). To visualize the clusters in a two-dimensional space, the Uniform Manifold Approximation and Projection (UMAP) algorithm was used and ClusterExplorer enabled 1) the assessment of each marker expression on the identified clusters and 2) the quantification of the clusters’ frequency in each treatment group. Finally, to quantitatively label the clusters, the Marker Enrichemnt Modeling (MEM)^52^ tool was used with a marker inclusion threshold set to 2. All the algorithms described were used as FlowJo plugins.

### Statistics

Data are expressed as mean ± standard error of the mean (SEM). Statistical analyses were conducted using GraphPad PRISM 10, with significance defined as *p ≤ 0.05, **p ≤ 0.01, and ***p ≤ 0.001. Comparisons between a single treatment group and the control group were evaluated using an unpaired two-tailed Student’s *t*-test. For analyses involving more than two experimental groups, one-way analysis of variance (ANOVA) was applied, followed by Tukey’s or Dunnett’s post hoc multiple comparison tests when statistical significance was observed (p < 0.05). Survival differences were assessed using the Mantel–Cox log-rank test.

## Data analysis

Analysis of flow cytometry was performed in FlowJo Version 10.10. Statistical analysis of data sets and data visualization was performed in Graphpad Prism 10.5.

## Author contributions

**M. Schork:** Conceptualization, data curation, formal analysis, investigation, methodology, project administration, visualization, validation, writing – original draft. **T. Torcellan:** Data curation, formal analysis, investigation, methodology, validation, visualization, writing – original draft, writing – review and editing. **I. Tsoukala:** Data curation, formal analysis, investigation, methodology, software, visualization. **A. Federico:** Data curation, formal analysis, investigation, methodology, software, visualization. **E. Diken:** Data curation, formal analysis, methodology, software, visualization. **O. Akilli:** Data curation, formal analysis, methodology, validation, visualization. **J. Gerbracht:** Data curation, formal analysis, methodology, software, visualization. **L. Kolb:** Methodology, supervision. **J. Zimmermann:** Data curation, formal analysis, investigation. **K. Wohde:** Investigation, visualization. **I. Beulshausen:** Investigation, visualization. **M. Suchan:** Data curation, formal analysis, methodology. **K. Khinvasara:** Formal analysis, software. **F. Beleggia:** Investigation, resources. **H. C. Reinhardt:** Funding acquisition, resources. **F. Lang:** Formal analysis, software, visualization. **B. Schrörs:** Formal analysis, methodology, Software, visualization, resources. **U. Sahin:** Funding acquisition, supervision. **M. Diken:** Funding acquisition, supervision. **A. Kübler:** Conceptualization, data curation, formal analysis, funding acquisition, methodology, project administration, supervision, visualization, writing - original draft, writing – review and editing.

All authors contributed to the revision of the manuscript, approved the final version, and consented to its publication.

## Acknowledgement

The authors thank Roman Thomas, Graziella Bosco and the whole CRC1399 of the DFG (grant no. 413326622) which supported parts of this study financially. H.C.R. was supported by the German Research Foundation (DFG) (SFB1399-grant no. 413326622, SFB1430 – grant no. 424228829, and SFB1530 – grant no. 455784452, RE 2246/17-1 – 553375105, SFB1752 – grant no. 549898653), the German Cancer Aid (1117240, 70113041, the Mildred Scheel Nachwuchszentrum Grant 70113307, the TACTIC consortium as part of the Preclinical Drug Development Program preCDD and an Excellence Funding Program grant, the TransOnk consortium ERASE), the German Ministry of Education and Research (BMBF e:Med Consortium InCa, grants 01ZX1901, 01ZX2201A and the DKTK grant BOOST-CAR-T) and the CANcer TARgeting (CANTAR) project NW21-062 “Netzwerke 2021” an initiative of the Ministry of Culture and Science of the State of North Rhine-Westphalia. The authors thank Sebastian Attig, Javad Assarzadeh and Kathrin Claude for their technical assistance in flow cytometry and cell sorting; Sonja Witzel and Melanie Grässl for support in cloning lentiviral vectors as well as vaccine constructs; Hanna Beyer and Mario Perkovic for assistance on lentivirus production; Alekhya Porapu, Viktoria Pitthan, Münteha Yilmaz, Michel Tenspolde, Elena Klein, Michelle Jäger, Anna Schmidt, Aninditha Muralidharan, Celine Bitter and Yvonne Feuchter for excellent in vivo and in vitro assistance; Nele Brüne and Diana Schneider for technical assistance on enzyme measurements; Jenny Schunke, Fulvia Vascotto, Karl-Heinz Tomaszowski and Nadja Salomon for scientific discussion; Aras Toker for critical feedback to the manuscript; David Weber, Thomas Bukur, Alina Henrichs, Feyza Durak, Aline Beicht and Lisa Vetterdietz for technical and scientific assistance on sequencing; Jeannette Rudzitis-Auth, Marianne Mathäs, Marleen Büchler-Schäff, Lotte Hofmann, Janne Bredehöft and Kurt Reifenberg for assistance on legal aspects regarding *in vivo* experiments. The authors want to especially acknowledge the assistance of Michael Kuske and Abderaouf Selmi in their role as direct supervisors of animal experiments as well as their scientific input.

## Data availability

The data supporting the findings of this study are available within the Article and its Supplementary and Extended Data files. Additional data are available from the corresponding author upon reasonable request. Sequencing data generated in this study will be deposited in an appropriate publicly accessible repository prior to final publication. Accession numbers and repository details will be provided in the published article.

## Materials availability

Materials generated in this study are available from the corresponding author upon reasonable request, subject to institutional and material transfer requirements.

## Artificial intelligence statement

During the preparation of this manuscript, the authors used AI tools for language refinement and editing to improve clarity and readability. All AI-assisted revisions were critically reviewed by the authors, who take full responsibility for the accuracy and integrity of the final manuscript.

