## Extended Data Figures for "Overcoming subtype-specific immune resistance in small cell lung cancer through RNA-based immunotherapy"

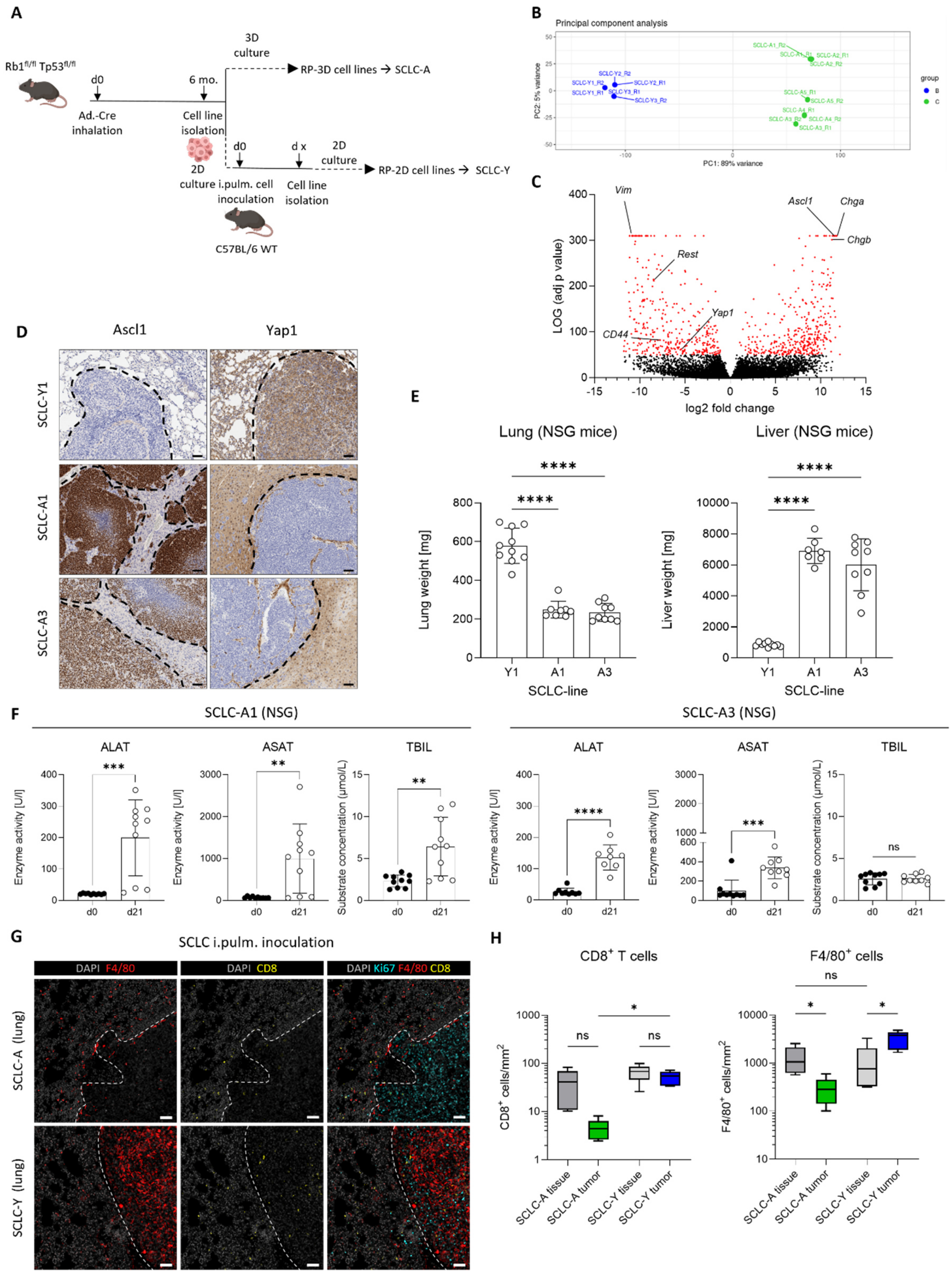

**Extended Data Figure 1: Murine SCLC cell line generation and characteristics. (A)** Rb1$^{fl/fl}$ Tp53$^{fl/fl}$ (RP)-C57BL/6 mice were intratracheally instilled with Adeno-Cre virus to initiate autochthonous tumorigenesis. Mice were sacrificed when relevant lung tumor burden was reached (ca. 6 months after inhalation), cell lines were generated from primary tumors from different mice. Cell lines were either cultured in 3D as described in Methods to generate SCLC-A subtype-reflecting cells or were cultured in 2D after another in vivo passage upon intrapulmonary injection and re-isolation. 2D cultured cells reflect the SCLC-Y subtype. Pictures partially created with Biorender (License and link in Extended Data Figure 3B) **(B)**

Principal component analysis from bulk RNA sequencing with technical duplicates per cell line. Blue dots represent 2D-cultured and green dots 3D-cultured cell lines. (**C**) Volcano plot showing differential gene expression from bulk RNA sequencing of SCLC cell lines. The x-axis represents the log2 fold change of gene expression between SCLC-Y and SCLC-A cell lines, the y-axis represents the log10 of the adjusted p-value (padj) indicating the statistical significance of the differential expression. Genes meeting the significance threshold ($log^{padj}$ = 50) are highlighted in red. Selected genes of interest are labeled. (**D**) Tumor-bearing mice (n = 5) at endpoint were histologically analyzed for subtype-defining Ascl1 and Yap1 (SCLC-A in liver, SCLC-Y in lung). Shown is one representative image, black dashed lines indicate tumor border, scale bar 50 µm. (**E**) Lung and liver weights of NSG mice upon intravenous injection of SCLC cell lines at individual endpoint (n = 10). Increased lung weight in SCLC-Y1 cell line showcases increased tumor burden in lung, whereas SCLC-A1 and A3 cell lines display increased tumor burden in liver as shown by increased liver weight. (**F**) Enzyme activity levels of alanin-aminotransferase (ALAT), aspartat-aminotransferase (ASAT) and substrate concentration of bilirubin (TBIL) in serum level of NSG mice at d 0 and d 21 after injection with either SCLC-A1 or SCLC-A3 as determined by clinical chemistry analysis. Significance was determined using a two-tailed t-test for paired samples. (**G**) SCLC-A lung tumor-bearing mice (n = 5) were histologically analyzed for infiltration of $CD8^+$ T cells, $F4/80^+$ cells and proliferation (Ki67). Shown is one representative image, dashed lines indicate tumor border. Scale bar = 50 µm (**H**) Quantification of $F4/80^+$ and $CD8^+$ cells in SCLC-A and SCLC-Y lung tumors as cells per $mm^2$. Analysis comprises tumor- and adjacent tissue. Statistical significance is indicated as follows: p < 0,05 = *; p < 0,01 = **; p < 0,001 = ***; p < 0,0001 = ****

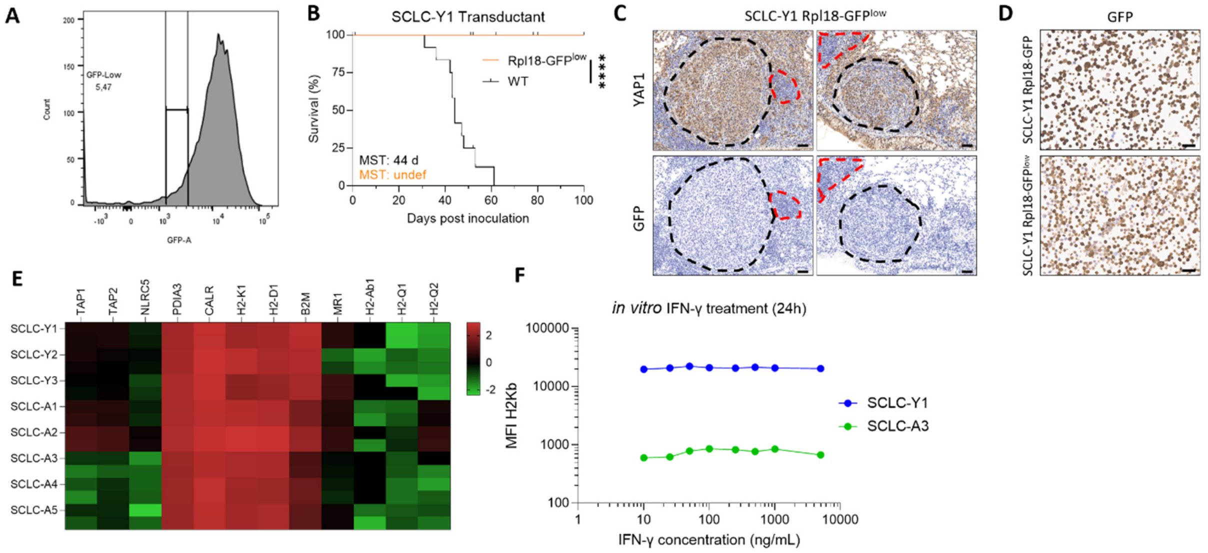


**Extended Data Figure 2: T cell recognition in SCLC-A and Y. (A)** SCLC-Y1 Rpl18-GFP cells were sorted based on GFP expression using the indicated gate, generating a SCLC-Y1 Rpl18-GFP low tumor cell line. (**B**) Engraftment of SCLC-Y1 Rpl18-GFP low sorted cell lines upon intravenous injection into C57BL/6 mice (n = 10) compared to WT SCLC-Y1 cells. Please note, the SCLC-Y1 wildtype survival is already shown in Fig. 1B. (**C**) Three survivor mice from SCLC-Y1 Rpl18-GFP low inoculated mice shown in B were analyzed histologically for tumor burden. One out of 3 mice showed residual tumor burden and was stained for YAP1 and GFP. Tumor tissue is indicated with a black dashed line, lymphatic aggregates with a red dashed line, scale bar 50 µm. (**D**) Cell pellet staining of GFP on SCLC-Y1 Rpl18-GFP and SCLC-Y1 Rpl18-GFP$^{low}$, scale bars = 50 µm (**E**) Expression level of genes involved in antigen presentation machinery from bulk RNA sequencing shown as $\log_{10}$ expression level of technical duplicates (**F**) Tumor cell lines were treated with indicated concentrations of recombinant IFN-γ for 24 hours, MHC-I (H2kb) expression was quantified by flow cytometry.

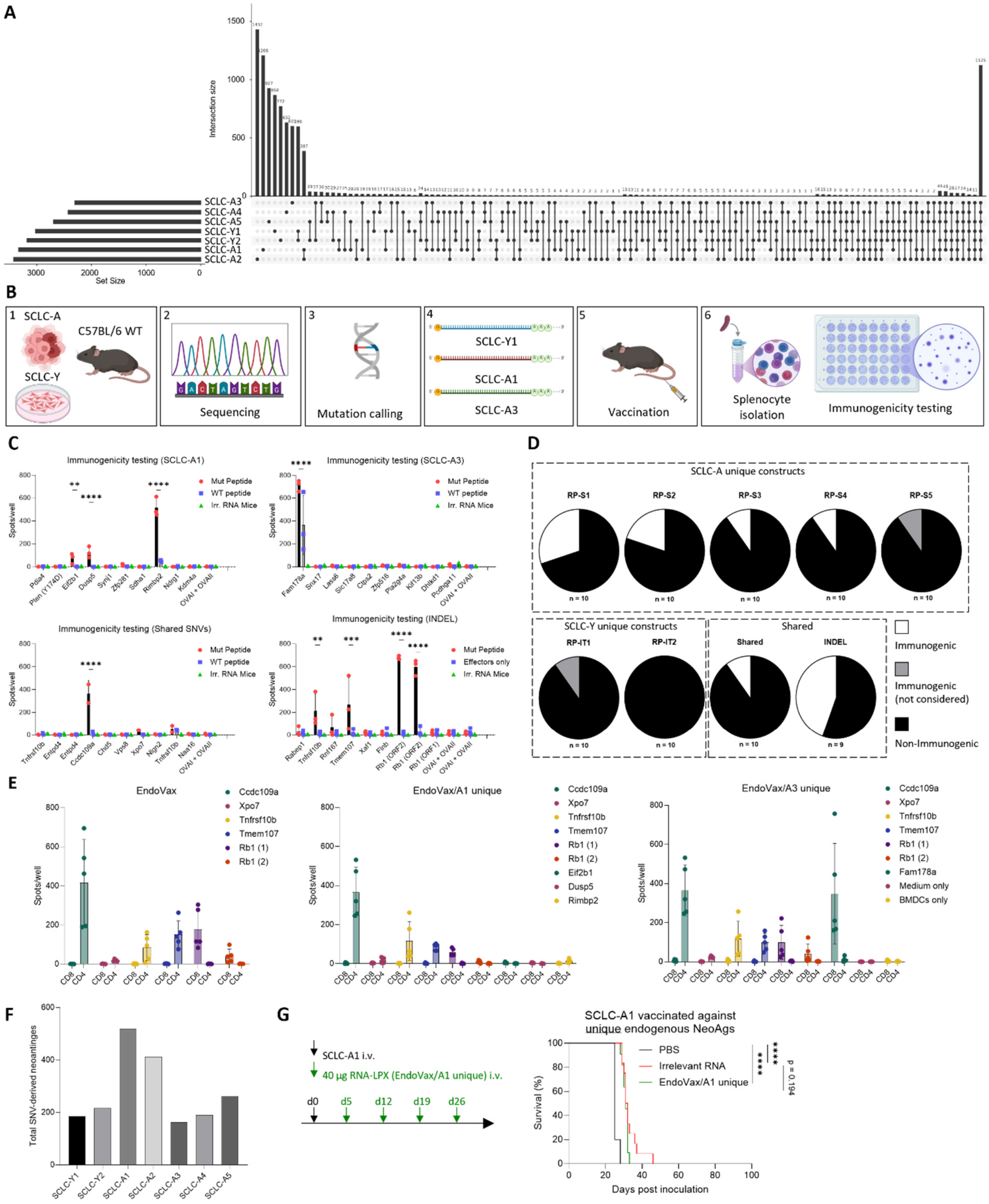


**Extended Data Figure 3: Tumor-specific antigen identification and validation. (A)** Intersection plot of somatic mutations in SCLC cell lines. The plot displays the number of mutations per cell line and intersection size in multiple samples/groups, highlighting both unique (individual, as shown by dots) and shared variants. (**B**) Tumor specific antigen prediction and validation pipeline. Briefly, SCLC cell lines were subjected to sequencing, and neoantigen candidates were identified using a computational prediction workflow. Neoantigen candidates were then cloned into expression vectors for in vitro transcription (IVT) RNA production and used to immunize naïve C57BL/6 mice. Following vaccination, $CD4^+$ and $CD8^+$ T cells were isolated from splenocytes and re-stimulated ex vivo with mutant candidate peptides or corresponding wild-type control peptides. Antigen-specific T-cell responses were assessed by IFN-γ ELISpot, enabling the identification of tumor-specific antigens by positive IFN-γ ELISpot (Created in BioRender. https://BioRender.com/s68lfhh). (**C**) Shown are IFN-γ spots per well after C57BL/6 immunization with candidate antigen RNA constructs that were either derived from SCLC-A1, SCLC-A3, shared SNVs or shared INDELs and ex vivo re-stimulation of splenocytes with mutated (antigen candidates) or wildtype peptides as well as irrelevant RNA-LPX (empty vector RNA) vaccinated mice. SCLC-Y1 harbored no considered immunogenic neoantigens. (**D**) Pie charts showing the proportion of validated immunogenic antigens among the 9-10 candidates tested per RNA construct. Non-considered immunogenic neoantigens include those, on which the WT peptide also elicits T cell responses. (**E**) Immunogenic antigens were re-cloned and validated in vivo. Naïve C57/BL6 mice (n = 5) were immunized with RNA-LPX, CD8 or CD4 T cells were isolated from splenocytes and re-stimulated with peptides ex vivo to determine, if the antigen elicit CD8 and/or CD4 responses. Tested were Endovax (Shared SNVs + InDel fusion) fused to respective unique neoantigens. (**F**) Total number of identified SNV-derived predicted neoantigen-candidates per cell line. (**G**) Vaccination of SCLC-A1 WT tumor bearing mice against identified unique neoantigens. No significant survival effect was achieved by neoantigen coding vs. irrelevant RNA, indicating no relevant T cell effect. Significance levels are calculated using log-rank (Mantle-Cox) test.

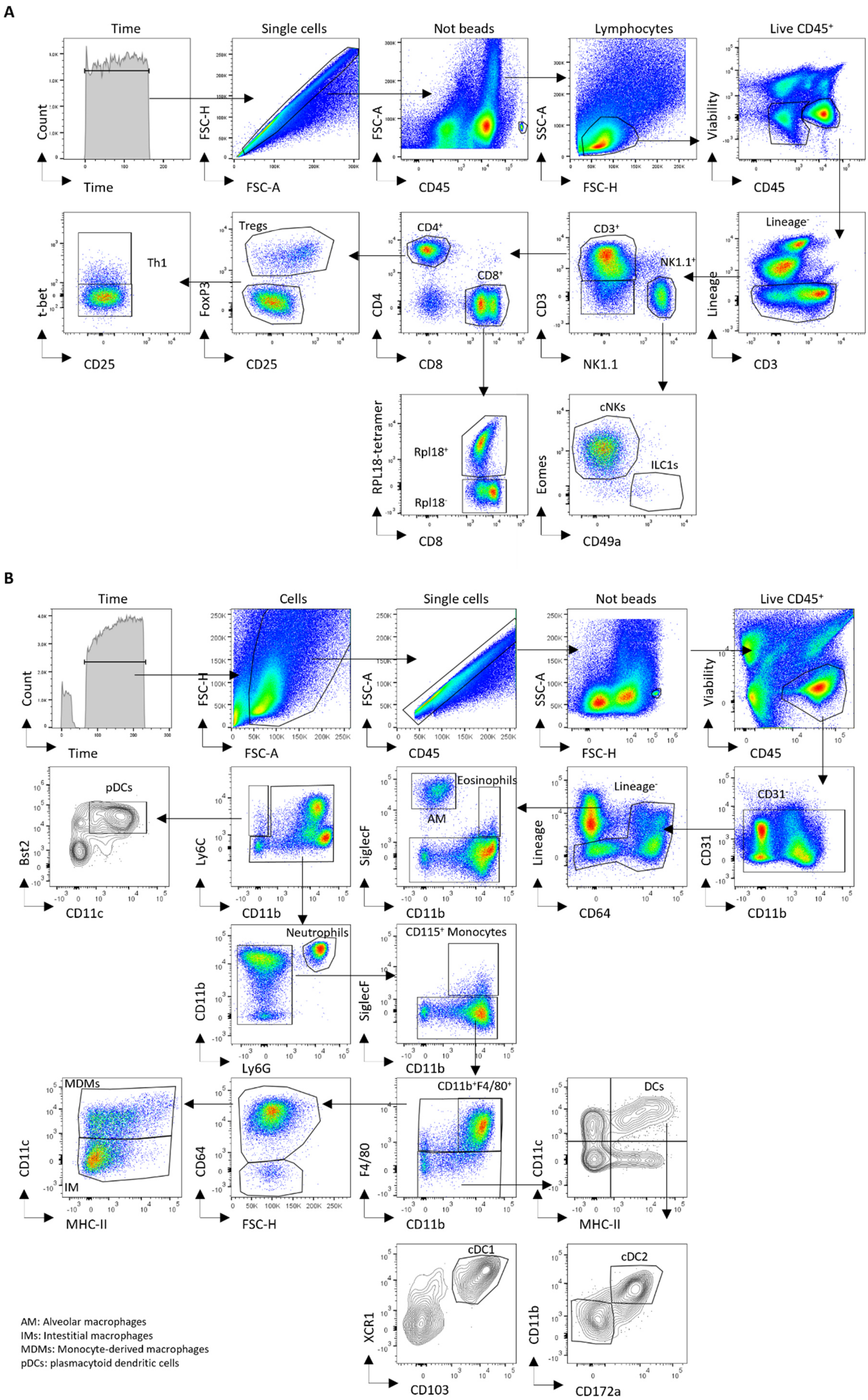


AM: Alveolar macrophages
IMs: Intestitial macrophages
MDMs: Monocyte-derived macrophages
pDCs: plasmacytoid dendritic cells

**Extended Data Figure 4: Gating strategy of myeloid subsets and lymphoid subsets in Experiment Fig 4 & 5.** (**A**) Gating strategy on lymphoid subsets shown on representative SCLC-A3 Rpl18-GFP tumor-bearing lungs after Rpl18/Endovax RNA vaccination (**B**) Gating strategy of myeloid subsets shown on representative lung samples harboring SCLC-A3 Rpl18-GFP tumors.

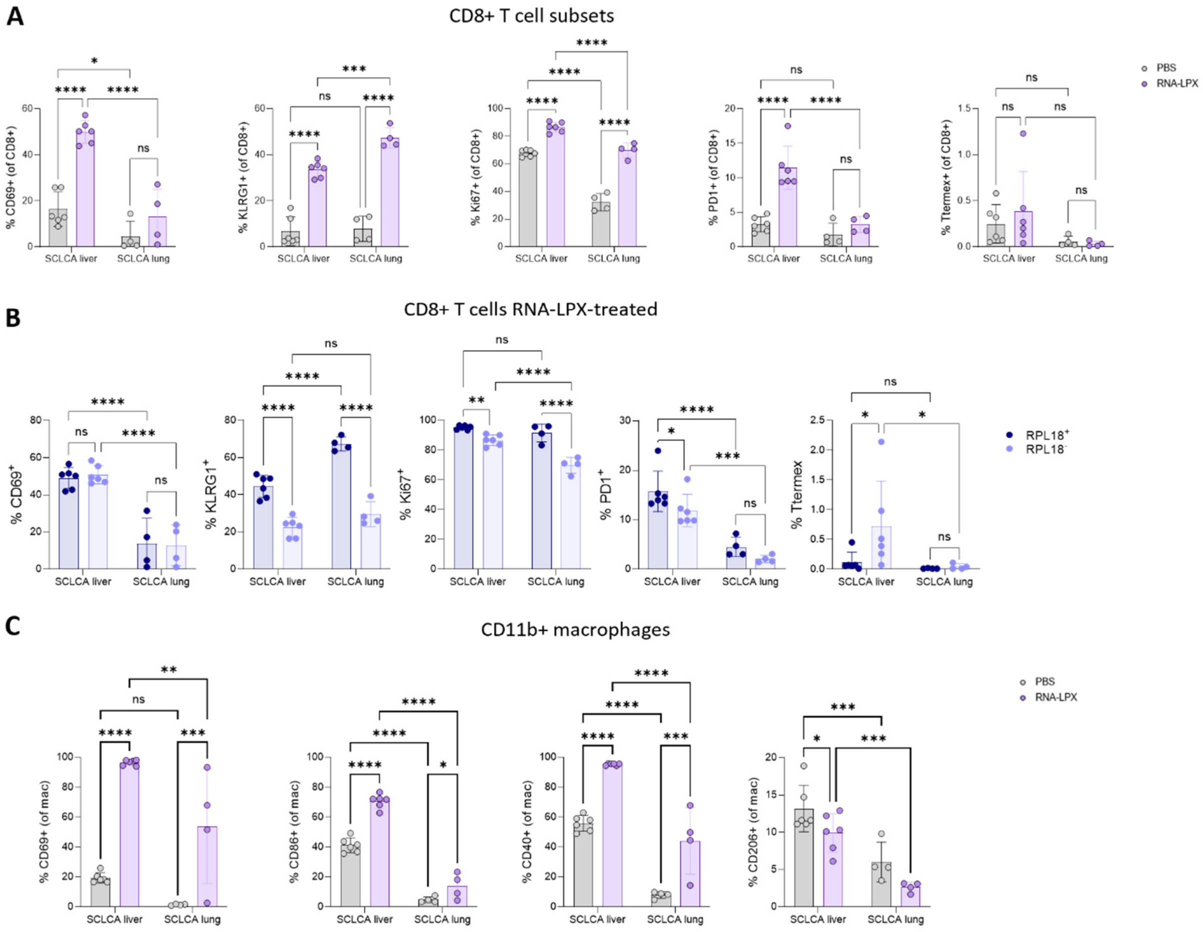


**Extended Data Figure 5: Flow cytometric analysis of SCLC-A tumor-bearing lungs and livers after RNA-LPX vaccination (A)** Expression of CD69, Klrg1, Ki67, PD-1 and terminal exhaustion markers (PD-$1^+$ CD$101^+$) on all CD$8^+$ cells defined per gating strategy in Extended Data Figure S4. Significancy levels were calculated using Two-Way ANOVA with Tukey's post-hoc correction (**B**) Expression of markers in (A) in direct comparison between SCLC-A tumor bearing organs in neoantigen-specific vs. non-neoantigen specific T cells. Statistical significance was calculated by Two-Way ANOVA with Tukey's post-hoc correction (**C**) CD69, CD86, CD40 and CD206 expression on CD11b$^+$ macrophages between lung and liver SCLC-A tumor bearing organs. Statistical significance was assessed by Two-Way ANOVA with Tukey's post-hoc test for multiple comparisons. Statistical significance is indicated as follows: $p < 0.05$ = *; $p < 0.01$ = **; $p < 0.001$ = ***; $p < 0.0001$ = ****

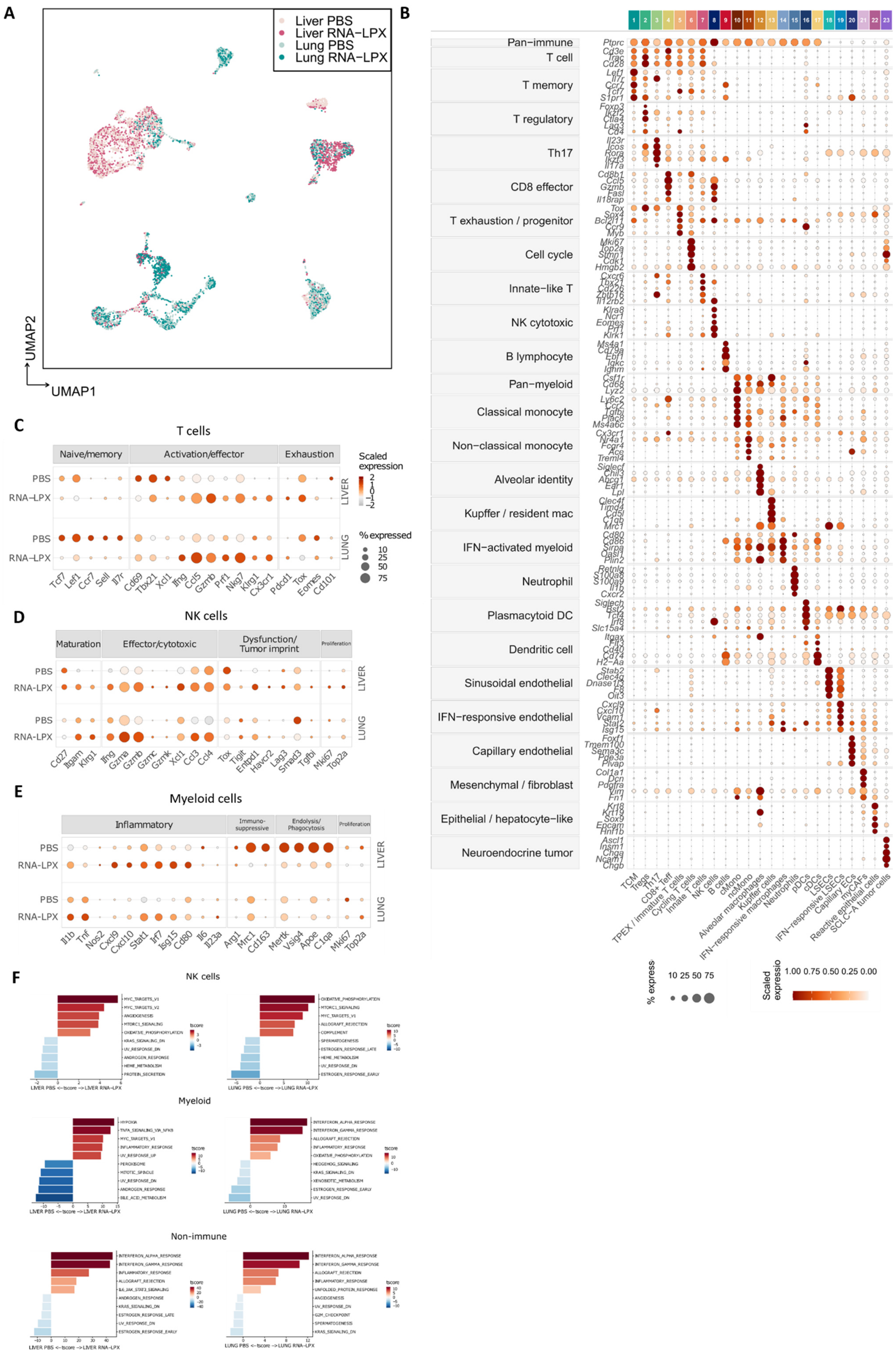

A
Liver PBS
Liver RNA−LPX
Lung PBS
Lung RNA−LPX
UMAP2
UMAP1
B
Pan−immune
T cell
T memory
T regulatory
Th17
CD8 effector
T exhaustion / progenitor
Cell cycle
Innate−like T
NK cytotoxic
B lymphocyte
Pan−myeloid
Classical monocyte
Non−classical monocyte
Alveolar identity
Kupffer / resident mac
IFN−activated myeloid
Neutrophil
Plasmacytoid DC
Dendritic cell
Sinusoidal endothelial
IFN−responsive endothelial
Capillary endothelial
Mesenchymal / fibroblast
Epithelial / hepatocyte−like
Neuroendocrine tumor
TCM
Tregs
Th17
CD8+ Teff
TPEX / immature T cells
Cycling T cells
Innate T cells
NK cells
B cells
cMono
ncMono
Alveolar macrophages
Kupffer cells
IFN−responsive macrophages
Neutrophils
pDCs
cDCs
LSECs
IFN−responsive LSECs
Capillary ECs
myCAFs
Reactive epithelial cells
SCLC−A tumor cells
% express
10 25 50 75
Scaled expressio
1.00 0.75 0.50 0.25 0.00
C
T cells
Naive/memory
Activation/effector
Exhaustion
Scaled expression
% expressed
PBS
RNA−LPX
LIVER
LUNG
D
NK cells
Maturation
Effector/cytotoxic
Dysfunction/ Tumor imprint
Proliferation
E
Myeloid cells
Inflammatory
Immuno-suppressive
Endolysis/ Phagocytosis
Proliferation
F
NK cells
Myeloid
Non-immune
LIVER PBS <−tscore −> LIVER RNA−LPX
LUNG PBS <−tscore −> LUNG RNA−LPX
tscore

**Extended Data Figure 6: snRNA-seq analysis of the multiome dataset** **(A)** UMAP map of analysis showing the cell distribution based on treatment groups. (**B**) Top differentially expressed genes on identified clusters. Colors indicate scaled expression and bubble size indicates expressing fraction (**C**) Gene expression patterns, across treatment and site groups, of T cells, (**D**) NK cells and (**E**) myeloid cell markers accounting for selected biological programs. (**F**) GO term analysis of indicated cell populations showing increased and decreased biological processes upon treatment

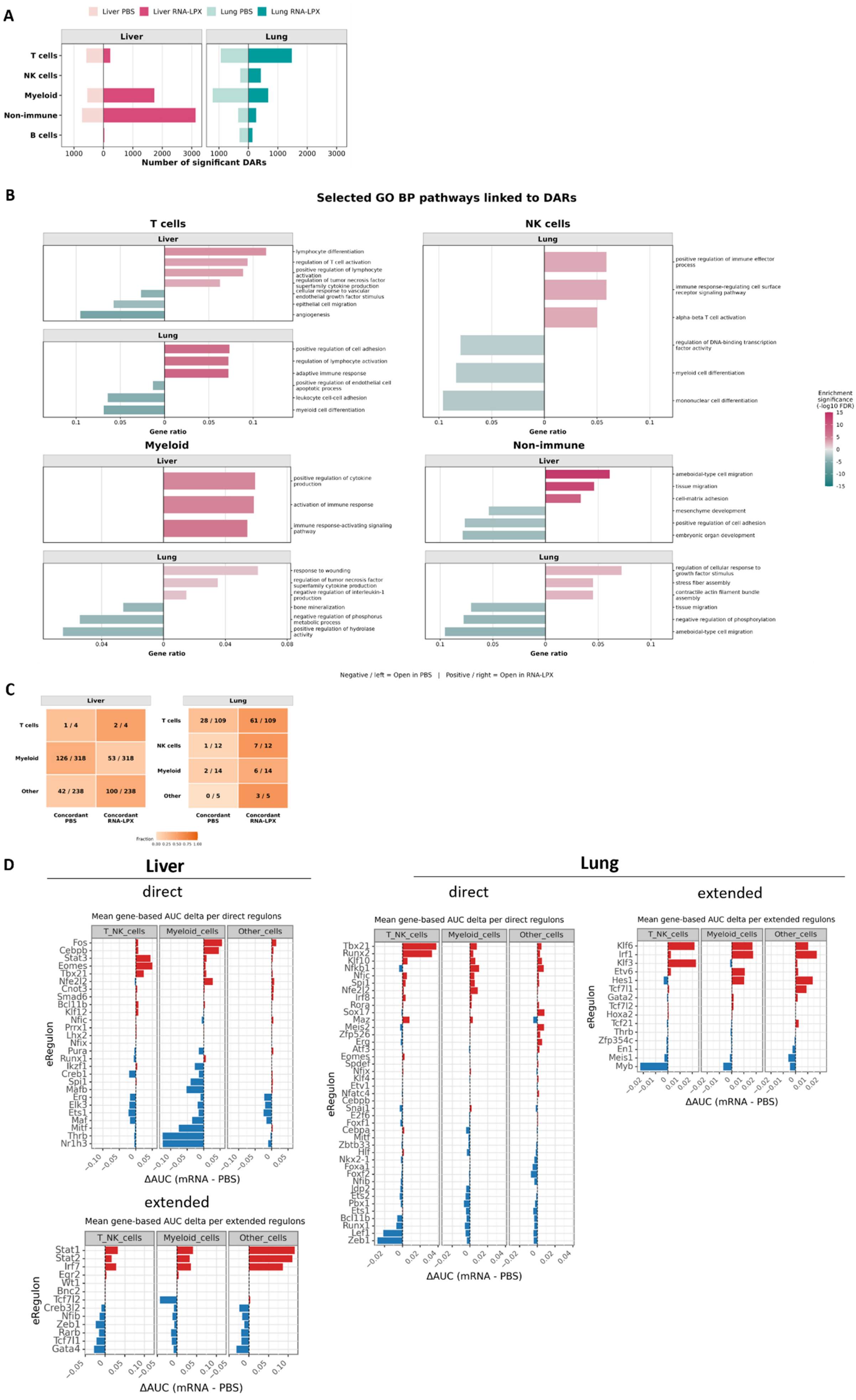
A
Liver PBS
Liver RNA-LPX
Lung PBS
Lung RNA-LPX
Liver
Lung
T cells
NK cells
Myeloid
Non-immune
B cells
Number of significant DARs
B
Selected GO BP pathways linked to DARs
T cells
NK cells
Myeloid
Non-immune
Gene ratio
Enrichment significance (-log10 FDR)
lymphocyte differentiation
regulation of T cell activation
positive regulation of lymphocyte activation
regulation of tumor necrosis factor superfamily cytokine production
cellular response to vascular endothelial growth factor stimulus
epithelial cell migration
angiogenesis
positive regulation of cell adhesion
regulation of lymphocyte activation
adaptive immune response
positive regulation of endothelial cell apoptotic process
leukocyte cell-cell adhesion
myeloid cell differentiation
positive regulation of immune effector process
immune response-regulating cell surface receptor signaling pathway
alpha-beta T cell activation
regulation of DNA-binding transcription factor activity
myeloid cell differentiation
mononuclear cell differentiation
positive regulation of cytokine production
activation of immune response
immune response-activating signaling pathway
response to wounding
regulation of tumor necrosis factor superfamily cytokine production
negative regulation of interleukin-1 production
bone mineralization
negative regulation of phosphorus metabolic process
positive regulation of hydrolase activity
ameboidal-type cell migration
tissue migration
cell-matrix adhesion
mesenchyme development
positive regulation of cell adhesion
embryonic organ development
regulation of cellular response to growth factor stimulus
stress fiber assembly
contractile actin filament bundle assembly
tissue migration
negative regulation of phosphorylation
ameboidal-type cell migration
Negative / left = Open in PBS | Positive / right = Open in RNA-LPX
C
Liver
T cells 1 / 4 2 / 4
Myeloid 126 / 318 53 / 318
Other 42 / 238 100 / 238
Lung
T cells 28 / 109 61 / 109
NK cells 1 / 12 7 / 12
Myeloid 2 / 14 6 / 14
Other 0 / 5 3 / 5
Concordant PBS
Concordant RNA-LPX
Fraction
D
Liver
Lung
direct
extended
Mean gene-based AUC delta per direct regulons
Mean gene-based AUC delta per extended regulons
T_NK_cells
Myeloid_cells
Other_cells
eRegulon
ΔAUC (mRNA - PBS)

**Extended Data Figure 7: Analysis of differentially accessible regions by snATAC-seq** (**A**) Barplot showing the number of significantly DARs upon treatment in indicated cell populations across lungs and livers. **(B)** Selected DAR-inferred GO biological pathways across different cell types and organs. (**C**) Number of genes found concordantly deregulated, at both ATAC and gene expression level, based on treatment,, cell type and tumor site. B cells are not shown because no genes met the RNA–ATAC concordance criteria. (**D**) Waterfall plot showing all identified eRegulons per cell subset and organ. Direct eRegulons show high-confidence motif-to-TF annotation, extended show predicted eRegulons based on lower motif-to-TF assignments such as orthology to humans or motif similarity without direct ChIP-Seq annotation as defined by SCENIC+.

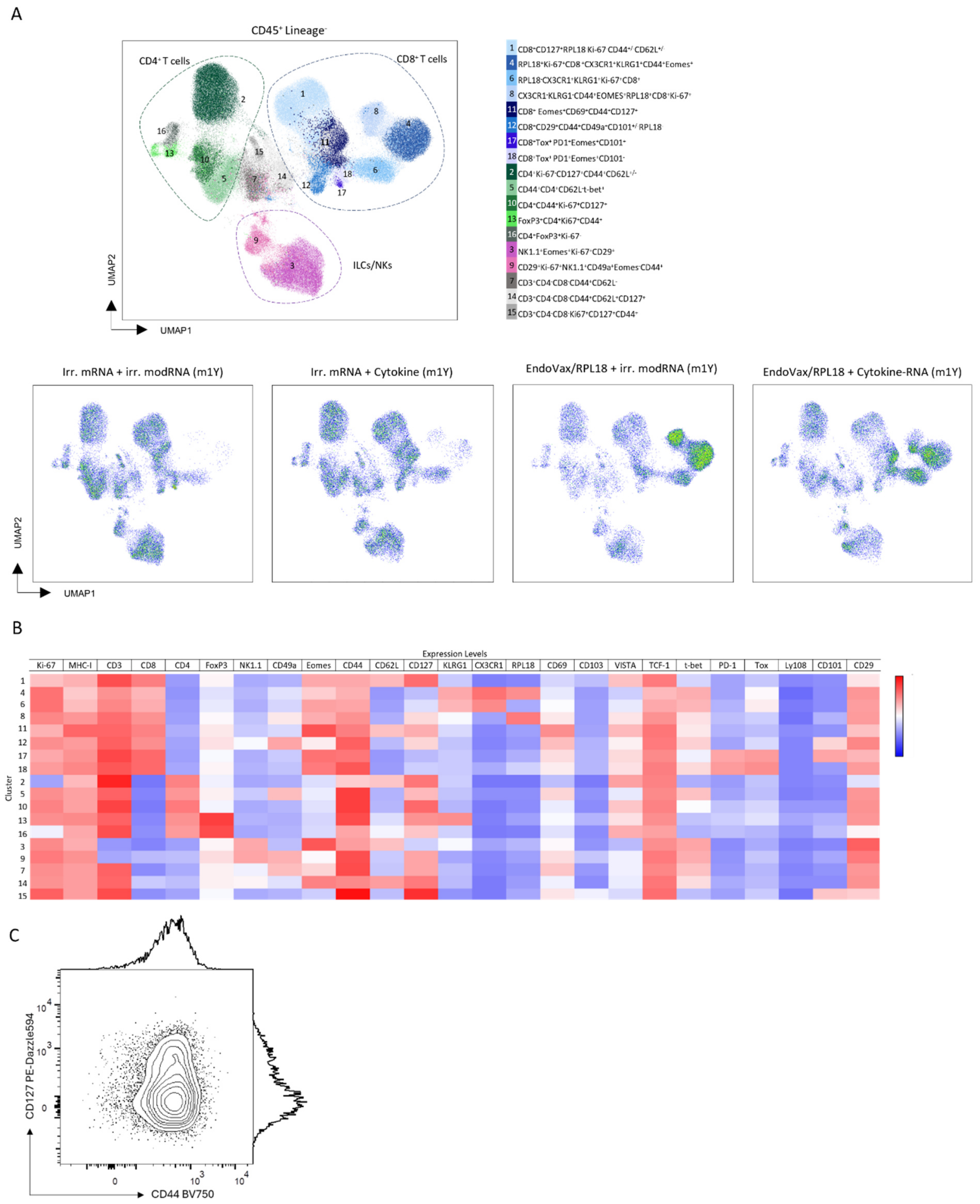


**Extended Data Figure 8: Cluster differences and identification on flow cytometry data.** (**A**) Dimensionality reduction analysis displayed in UMAP with clusters identified by X-Shift of liver samples from mice treated with NeoAg-encoding mRNA(U), Cytokine-encoding mRNA (m1Y), both or irrelevant RNA. Pseudocolor UMAP plots for each treatment group extracted from the global UMAP on the left **(B)** Heatmap displaying the relative intensity of each parameter for the clusters identified by X-Shift from four concatenated treatment groups. Color scaling uses full range of consensus transform (**C**) Dot plot of cluster 8, showing expression of CD44 and CD127.

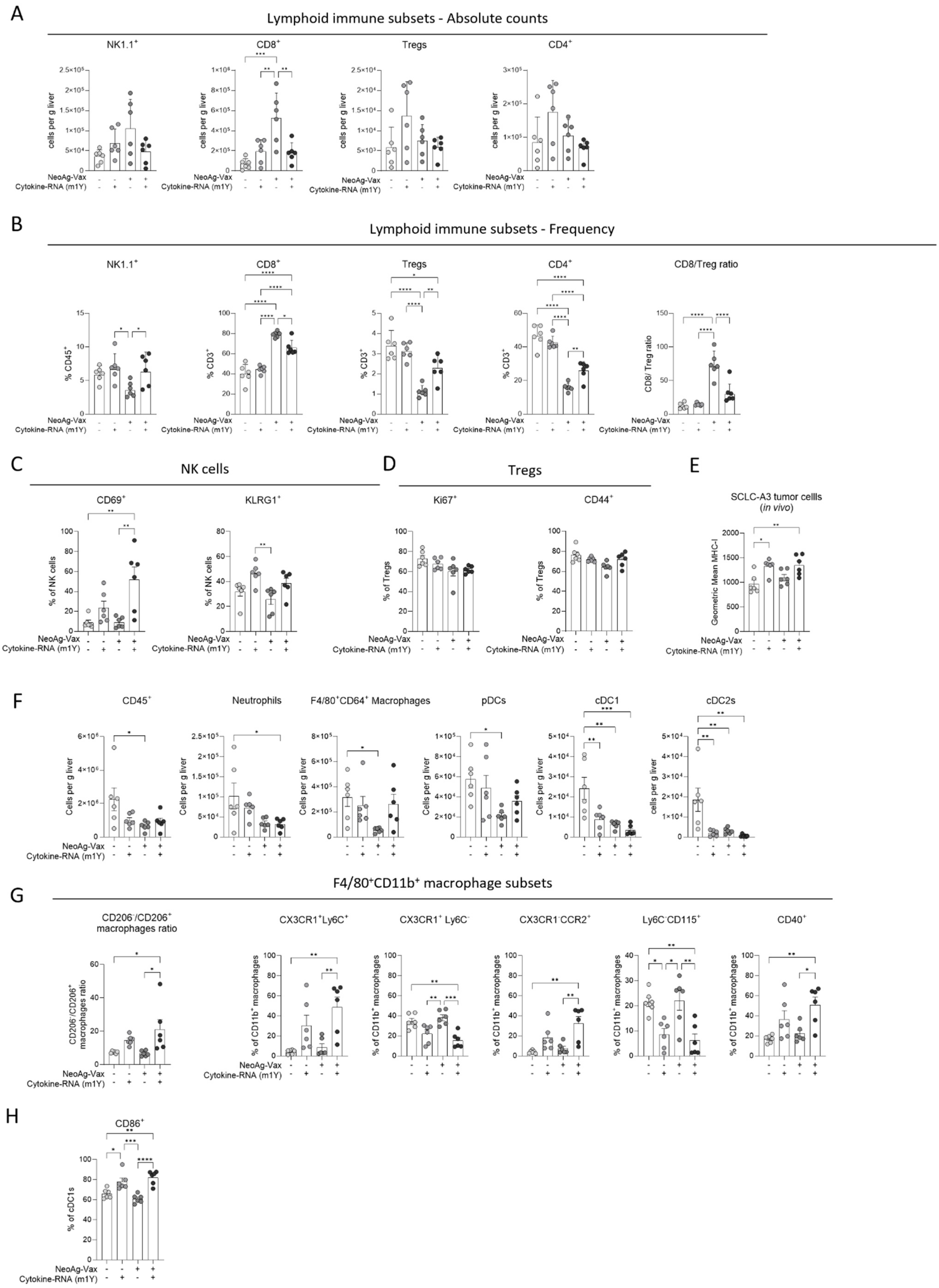
A
Lymphoid immune subsets - Absolute counts
NK1.1+
CD8+
Tregs
CD4+
cells per g liver
NeoAg-Vax
Cytokine-RNA (m1Y)
B
Lymphoid immune subsets - Frequency
NK1.1+
CD8+
Tregs
CD4+
CD8/Treg ratio
% CD45+
% CD3+
CD8/ Treg ratio
C
NK cells
CD69+
KLRG1+
% of NK cells
D
Tregs
Ki67+
CD44+
% of Tregs
E
SCLC-A3 tumor celils
(in vivo)
Geometric Mean MHC-I
F
CD45+
Neutrophils
F4/80+CD64+ Macrophages
pDCs
cDC1
cDC2s
Cells per g liver
G
F4/80+CD11b+ macrophage subsets
CD206-/CD206+ macrophages ratio
CX3CR1+Ly6C+
CX3CR1+ Ly6C-
CX3CR1-CCR2+
Ly6C-CD115+
CD40+
% of CD11b+ macrophages
H
CD86+
% of cDC1s

**Extended Data Figure S9: Characterization of lymphoid and myeloid subsets in the tumor immune microenvironment after RNA-LPX and RNA-encoded cytokine therapy in SCLC-A**. Flow cytometric analysis derived from the experiment shown in Figure 6, assessing SCLC-A tumor-bearing livers after RNA-LPX vaccination in combination with RNA-encoded cytokines. (**A**) Absolute counts per organ weight in livers based on treatment. (**B**) Frequencies of lymphoid subsets, (**C**) NK cell phenotype or (**D**) Tregs. (**E**) MHC-I expression on SCLC-A4 tumor cells in vivo. Significance was determined by one-way ANOVA with Tukey's post-hoc test for multiple comparisons. (**F**) Absolute counts per organ weight in livers based on treatment. (**G)** Frequencies of myeloid subsets, (**H**) CD86 frequency on cDC1. Significance was determined by one-way ANOVA with Tukey's post-hoc test for multiple comparisons. Statistical significance is indicated as follows: $p < 0.05$ = *; $p < 0.01$ = **; $p < 0.001$ = ***; $p < 0.0001$ = ****
